# Heterochromatin boundaries maintain centromere position, size and number

**DOI:** 10.1101/2025.02.03.635667

**Authors:** Ben L Carty, Danilo Dubocanin, Marina Murillo-Pineda, Marie Dumont, Emilia Volpe, Pawel Mikulski, Julia Humes, Oliver Whittingham, Daniele Fachinetti, Simona Giunta, Nicolas Altemose, Lars E.T. Jansen

## Abstract

Centromeres are chromosomal loci that ensure proper chromosome segregation by providing a platform for kinetochore assembly and spindle force transduction during cell division. Human centromeres are defined primarily by a unique chromatin domain featuring the histone H3 variant, Centromere Protein A (CENP-A), that marks a single active centromere locus per chromosome. CENP-A chromatin typically occupies a small subregion of low DNA methylation within multi-megabase arrays of hypermethylated alpha-satellite repeats and constitutive pericentric heterochromatin. However, the mechanisms defining and maintaining precise centromere position and domain size, and the role of the underlying alpha satellite DNA sequence, are poorly characterised. Using an experimentally-induced neocentromere in RPE1 cells, we discovered that the SUV39H1 and H2 methyltransferases tri-methylate H3K9 at neocentromere boundaries to maintain CENP-A domain size independent of DNA methylation or satellite sequences. Furthermore, we found that the CENP-A domain at canonical alpha-satellite-based centromeres is characterized by local depletion of H3K9me3-mediated heterochromatin, coinciding with the DNA methylation dip region. We identified the SETDB1 methyltransferase as key to maintaining H3K9me3 within flanking active higher order alpha satellite arrays while SUV39s and SUZ12 contribute to globally heterochromatinize both alpha satellites and neighbouring repeats. Loss of this heterochromatin boundary results in the progressive expansion of the primary CENP-A domain, erosion of DNA methylation, and the nucleation of new centromeres across alpha satellite sequences. Our study identifies the functional specialization of different H3K9 methyltransferases across centromeric and pericentric domains, crucial for maintaining centromere domain size and number.

## Introduction

Centromeres form the primary constriction of mitotic chromosomes, the chromosomal loci that drive chromosome segregation during mitosis and meiosis^1,2^. Despite its central role in cell division, how human centromere position and size are specified and how specifically one centromere per chromosome is maintained remains poorly understood. Human centromeres are characterised by multi-megabase arrays of AT-rich alpha satellite DNA^3,4^. Intriguingly, these sequences, while abundant, are not strictly required for centromere function. Instead, centromere position is, to a large extent, defined epigenetically in many eukaryotes, particularly in humans, and can be uncoupled from alpha satellite sequences. The key centromere-defining element is the histone H3 variant Centromere Protein A (CENP-A), which is sufficient to induce *de novo* centromere formation and appears as a central node in a self-propagating feedback loop that maintains centromere position^5–7^.

The dispensable nature of alpha-satellite DNA is further evident from the discovery of spontaneously formed centromeres at naïve loci, for instance in human patients^8–12^. These neocentromeres epitomize the epigenetic nature of centromeres, nucleating at sites devoid of alphoid DNA. Once formed, these neocentromeres are functional and propagated indefinitely on a human chromosome and, in certain cases, are even transgenerationally inherited^8,12,13^. Moreover, model systems such as human, yeasts, and chicken cells have been employed to experimentally induce neocentromere formation^14–18^. This can be achieved by deleting the endogenous centromere locus, thereby selecting for neocentromere seeding events to rescue chromosome stability. This approach allows us to isolate and monitor the evolution of spontaneously formed centromeres, and it provides an opportunity to explore the genetic and epigenetic contributions to centromere seeding and inheritance.

Centromeres are propagated through replenishment of centromeric chromatin in early G1-phase of each cell cycle^19–21^. New CENP-A histones are assembled onto chromatin in a self-propagating manner^6^, followed by replicative dilution in S-phase upon re-assembly onto each sister chromatid^21,22^. The self-templated, epigenetic feedback that maintains CENP-A position is predicted to generate variation in the precise positioning of the centromere due to the dynamic nature of histone recycling during DNA replication and transcription^23^. Indeed, centromere drift has been observed in evolutionary new centromeres of donkeys, as well as chicken Z chromosomes^24,25^. However, despite this local drift, vertebrate centromeres remain restricted to their broader chromosomal locus even if CENP-A is temporarily removed^26^, and their size remains restricted to ∼100-500 kilobases at both canonical and neocentromeres ^3,18,27,28^. The tightly constrained CENP-A domain is somewhat surprising as such “islands” of CENP-A typically reside in megabase-sized domains of homogenous alphoid repeats, indicating the existence of non-sequence-defined centromere boundaries.

One potential boundary is DNA methylation, specifically 5mC. Endogenous human centromeres occur within arrays of alpha satellite Higher Order Repeat (HORs) that generally have hypermethylated DNA, except for a distinct site of hypomethylation coinciding with strong enrichment of CENP-A chromatin, termed Centromere Dip Regions (CDRs)^3,29^. In addition, constitutive pericentromeric heterochromatin which flanks the centromere locus has been hypothesized to maintain centromere position^30^. This heterochromatin is marked by histone H3 lysine 9 trimethylation (H3K9me3), mediated by SUV39 methyltransferases, and it often spans other human satellite DNA families in pericentromeric regions, such as the classical Human Satellites 1-3 (HSats)^31,32^. H3K9me3 has been also observed to at least partially overlap with centromeric repeats^33–37^, and has recently been mapped using long-read sequencing approaches within active alpha satellite HOR arrays^38^, further indicating that the chromatin environment at and around the CENP-A locus may be of functional significance. However, whether heterochromatin or DNA methylation forms a true functional boundary and how this is regulated remain open questions. In this study, we have identified the enzymes that functionally maintain heterochromatin within centromere domains and are crucial to maintaining CENP-A domain position, size, and number.

## Results

### CENP-A domain at a nascent human neocentromere drifts while maintaining overall size

We previously developed a CRISPR-based neocentromere seeding system on chromosome 4 of human RPE1 cells to isolate and characterise *de novo* human neocentromere formation in an experimental environment (Fig 1A)^18^. This approach identified Neo4p13 – a 90 kb neocentromere that formed at a gene-poor region, within high levels of H3K9me3 constitutive heterochromatin^18^, analogous to canonical centromeres, offering a unique opportunity to probe its role in maintaining centromere position and size, independent of satellite sequences. To understand the positional stability of a newly seeded human neocentromere across cell division cycles, we employed Neo4p13 in an evolutionarily time-stamped laboratory setting to assess its mitotic stability during long-term passaging (Fig 1B). We took an early Neo4p13 isolate and proliferated three independent populations for 100 days. Then, from each independently evolved pool we randomly isolated three clonal subpopulations and assessed centromere position using CENP-A CUT&RUN (Fig 1C). From early passage (EP, red) cells to Day 100 (grey), we observed a gradual centromere drift over time (Fig 1D and E), consistent with previous studies in evolutionary new centromeres. Strikingly, however, the overall size of the Neo4p13 locus remained relatively tight at approximately 90-100 kb even as this centromere moved (Fig 1D and E). As the CENP-A domain drifts in one direction, CENP-A nucleosomes are depleted from the trailing end of the domain (e.g. Fig 1D Clones 1.2 and 3.2). These observations suggest that mechanisms exist to tightly maintain centromere boundaries and overall size of this domain.

**Figure 1:**
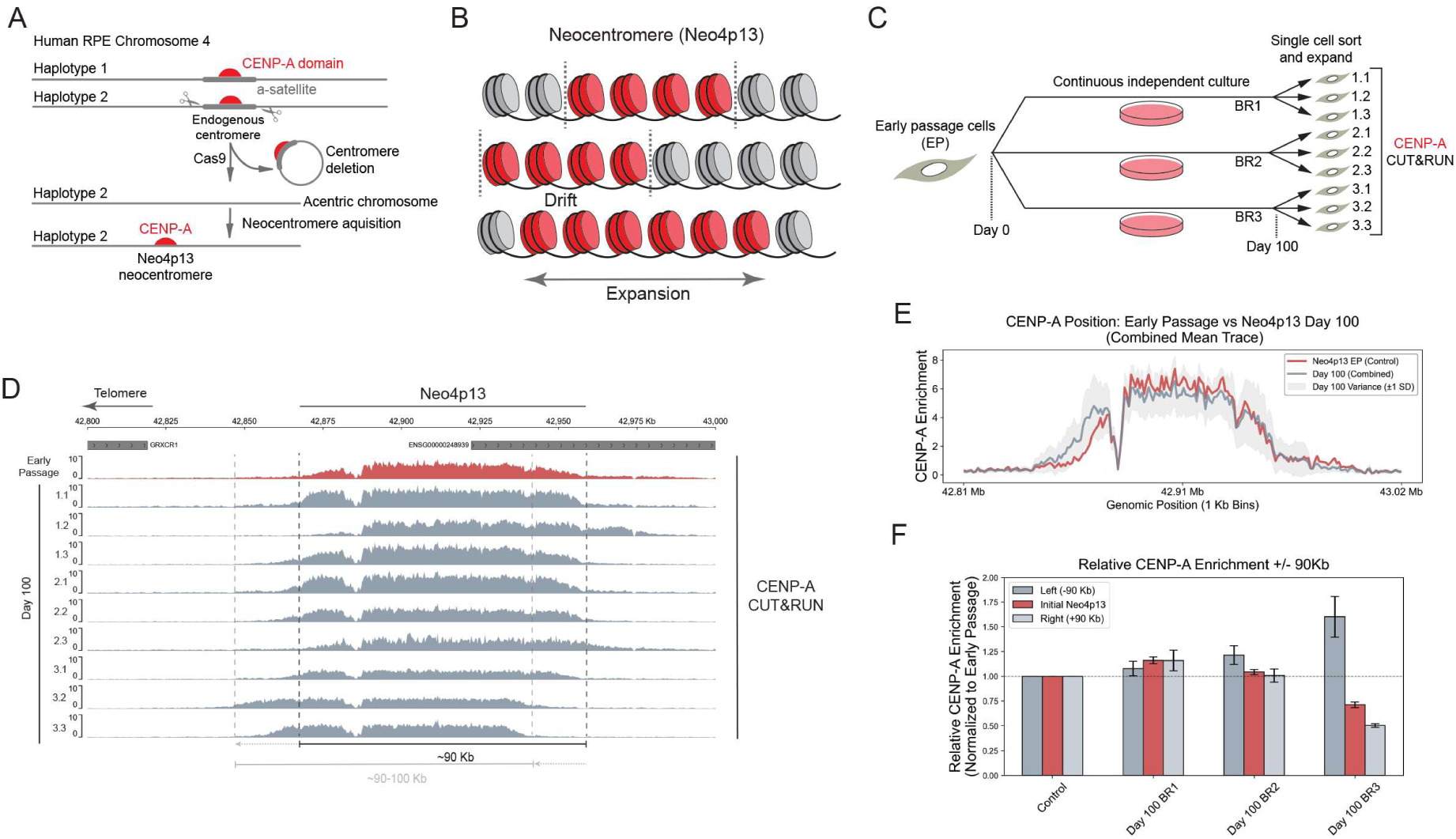
Long term Inheritance and size control of a human neocentromere. **(A)** Schematic of the generation of RPE1-Neo4p13, as previously described^18^. **(B)** Schematic predictions of centromere inheritance behaviour over time – centromeres may stay stationary, move left/right with tight size and boundary control, or gradually expand over time. **(C)** Schematic of long-term culture assay to monitor inheritance of Neo4p13 – culturing early passage Neo4p13 for 100 days, followed by single-cell sorting and CENP-A CUT&RUN (BR, Biological Replicate). **(D)** CENP-A CUT&RUN of early passage Neo4p13 (red) versus 100-day clones (grey). Black dotted lines represent early passage centromere position. Light grey dotted line represents position of clone 3.2 as an example centromere shift. **(E)** Quantitation of CENP-A enrichment across the genomic region 42.8–43.0 Mb (Chr 4 Hap 2) represented in 1 kb bins. Red trace represents the control sample (Neo4p13 EP), highlighting early passage CENP-A enrichment. The grey trace represent the combined mean enrichment for three groups of Day 100 clones: BR1, BR2, and BR3, with Day 100 +/− 1 standard deviation (SD, light grey) **(F)** Relative CENP-A enrichment in three genomic regions: the initial Neo4p13 locus (defined as 90 Kb window 42.868–42.958 Mb) and immediate flanking regions, 90 Kb upstream (“Left (−90 Kb)”) and downstream (“Right (+90 Kb)”). Enrichment values are normalized to the early passage control sample, represented by a baseline of 1 (dashed line). Each group of 100-day clones (BR1, BR2, and BR3) is displayed with bars representing relative enrichment in each region. Whiskers indicate the standard deviation (SD) across three individual clonal populations within each BR. Bar colours highlight the initial Neo4p13 locus (red, opaque) and the outside regions (greys).

### Seeding of Neo4p13 induces local depletion of 5mC and H3K9me3

To understand how neocentromere position is maintained, we mapped the local chromatin environment surrounding the CENP-A domain. The Neo4p13-RPE1 cells are diploid, effectively carrying two distinct centromere epialleles, one at the canonical centromeric alpha satellites of Chromosome 4 and one at 4p13 (Fig 1A). To accurately map chromatin features selectively at the neocentromere allele, we employed Oxford Nanopore (ONT)-based ultra-long-read DiMelo-seq^38^, in combination with the cell line-specific RPE1 reference genome^39^. This allows us to map single adenine-methylated DNA molecules in a haplotype-specific manner (Fig 2A). Neo4p13 maps to chromosome 4 haplotype 2 (Fig 2B, C), indicated by the presence of a CENP-A-directed 6mA peak at p13 on Hap2 specifically. Consistent with our previous observations using ChIP-seq^18^, this locus is enriched with H3K9me3 on both haplotypes. Interestingly, seeding of the CENP-A domain resulted in the selective loss of H3K9 trimethylation specifically at the neocentromere haplotype (Fig 2C and F).

**Figure 2:**
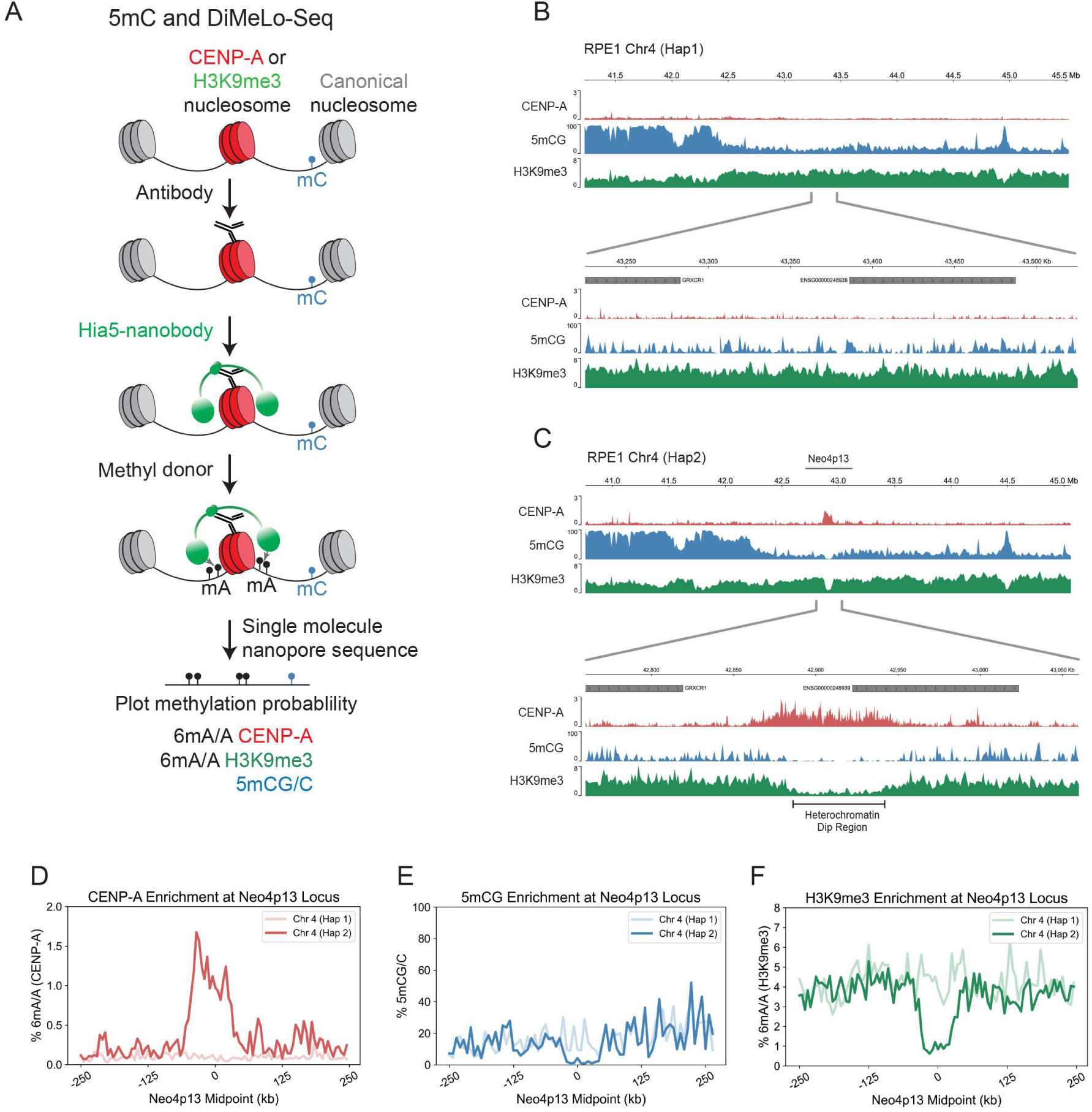
Neo4p13 occupies a distinct H3K9me3-mediated heterochromatin dip on Chr 4 Haplotype 2. **(A)** Schematic of the workflow of DiMelo-Seq and mapping to the diploid RPE1 genome. **(B and C)** Enrichment of CENP-A (red), H3K9me3 (green) and 5mCG (blue) at the genomic locus of Neo4p13, mapped to RPE1 chromosome 4 haplotype 1 **(B)** and chromosome 4 haplotype 2 **(C)**. Enrichment of CENP-A and dips in H3K9me3 and 5mCG are specific to chr4 hap 2. **(D)** CENP-A enrichment (% 6mA/A, red), **(E)** % 5mCG/C (blue), **(F)** H3K9me3 (% 6mA/A, green) at the Neo4p13 locus for two haplotypes: Chr4 (Hap1) and Chr4 (Hap2). Data is aggregated across a 500 kb region centred at the Neo4p13 coordinates, with flanking regions extending 250 kb on either side. Mean enrichment values are calculated in 100 evenly spaced bins per region. The x-axis represents the genomic position relative to the midpoint (in kilobases, kb), while the y-axis indicates the % 6mA/A or 5mCG/C.

Canonical alpha-satellite-based centromeres are characterised by distinct dips in 5mCG methylation at regions with high CENP-A occupancy^3,29,40^. Using DiMeLo-seq, we can directly measure endogenous 5mCG on the same single molecules on which we measure exogenous 6mA deposited around H3K9me3 or CENP-A. Using this approach, we found that the neocentromere arose in a broad domain of generally low DNA methylation (Fig 2B, C and E). Strikingly, the formation of the CENP-A domain induced the removal of the remaining 5mCG methylation, specifically on the CENP-A containing haplotype (Fig 2B, C and E). While we cannot directly distinguish between cause and consequence, our data on a newly formed neocentromere suggest that CENP-A seeding can specifically drive local CpG demethylation over time (Fig 2C-E).

Further, we analysed sequence features within the neocentromere domain. Previously, we determined that the overall AT-content of the 4p13 neocentromere region is roughly equal to that of the chromosome overall with only a slightly elevated AT content^18^. Here, we analysed the distribution of transposable elements (TEs) that have previously been implicated in centromere function, although the roles of TEs have been elusive^41^. Using the diploid RPE1 genome^39^, we characterised the TE distribution using RepeatMasker^42^. We compared Long terminal Repeats (LTRs), Long Interspersed Nuclear Elements (LINEs) and Short Interspersed Nuclear Elements (SINEs) across the neocentromere-containing chr4 (hap 2) (Fig S1A). As expected, all TEs are depleted from the canonical centromere, made up of satellites. Compared to q- and p- arms of the neocentromere-containing chromosome, the 4p13 neocentromere locus features a higher number of LTRs versus the chromosomal averages for Chr4 (Hap2) (Fig S1A and C). In contrast, LINEs and SINEs are closer to the genome average (Fig S1A-C). Importantly, while LTR numbers at 4p13 are elevated, many other regions across the chromosome feature higher numbers of all three TE classes (Fig S1A), making TEs unlikely to be defining features of this neocentromere.

### Heterochromatin at Neo4p13 forms functional centromere boundaries maintaining CENP-A domain size

Given the maintenance of size control of the 4p13 neocentromere that we observed in long-term culture, we hypothesised that adjacent H3K9me3-marked heterochromatin forms an effective boundary, preventing centromere expansion (Fig 3A). We predicted that the H3K9me3 methyltransferases SUV39H1 and SUV39H2 may be responsible for maintaining the heterochromatin domain at 4p13 as these are known to mark pericentric heterochromatin at canonical centromeres^43–45^. Furthermore, we also considered the H3K9me3 methyltransferase SETDB1 as a putative candidate, as this enzyme is known to have roles distinct from SUV39 enzymes, particularly in transposable element silencing^45–47^.

**Figure 3:**
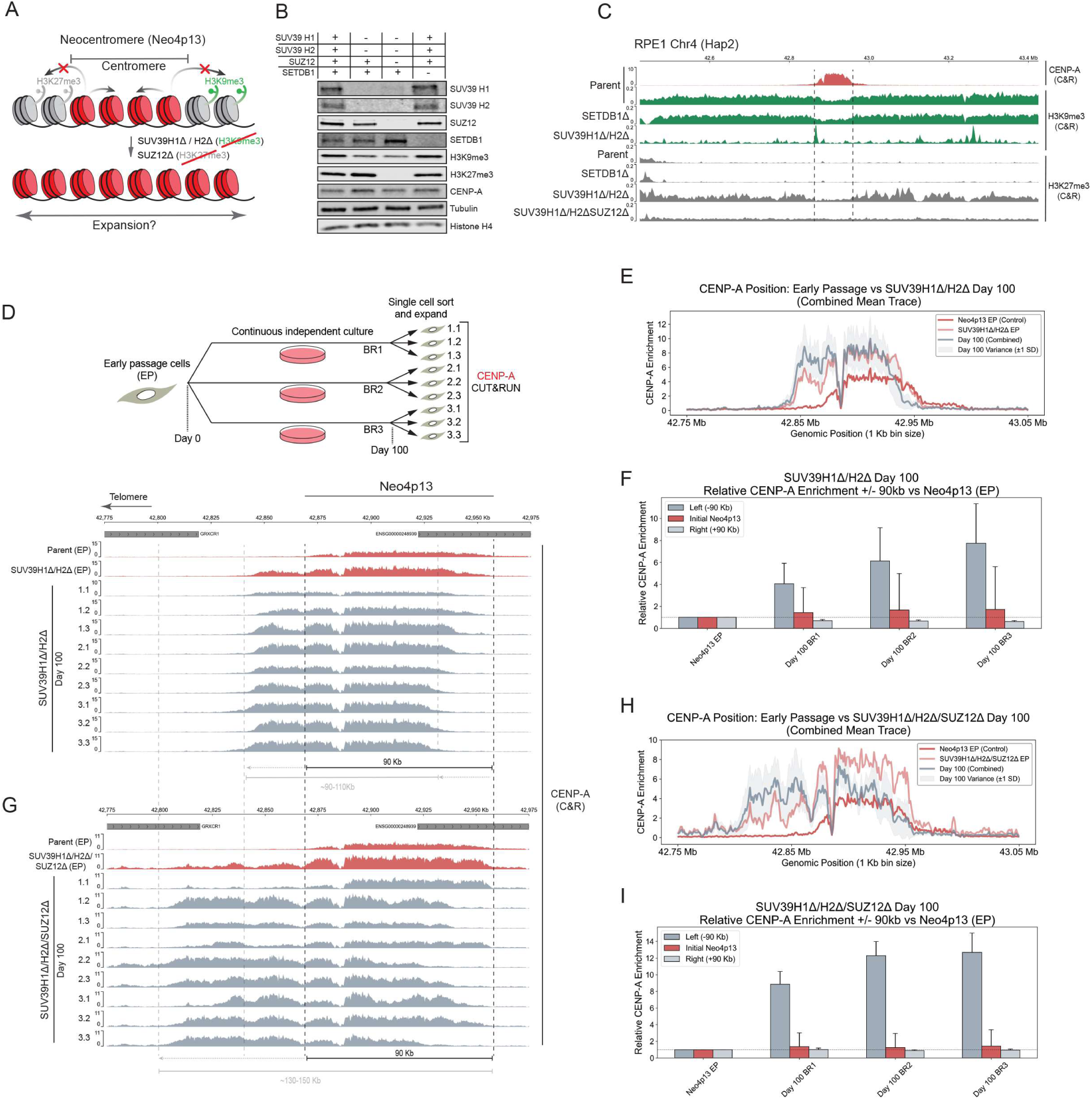
Heterochromatin forms functional boundaries and maintains tight size control of the Neo4p13 locus. **(A)** Model of the hypothetical role for heterochromatin as a functional boundary to maintain size control of the neocentromere. **(B)** Immunoblots of SUV39H1, SUV39H2 and Suz12 deletion lines, as single, double or triple knockouts, as well as SETDB1 single knockouts probed for indicated proteins and histone modifications. Histone H4 and α**-**tubulin serve as loading controls. **(C)** CUT&RUN of CENP-A (red), H3K9me3 (green) and H3K27me3 (dark grey) of indicated genotypes, across the broader Neo4p13 genomic locus. Dashed line represents the Neo4p13 centromere boundaries. **(D)** CENP-A CUT&RUN of early passage parental Neo4p13 (red) versus SUV39H1Δ/H2Δ (red) EP clones (red) versus corresponding 100-day clones (grey). Black dotted lines represent early passage centromere position. **(E)** Line graph of CENP-A enrichment across the genomic region 42.8–43.0 Mb (Chr 4 Hap 2) comparing parental Neo4p13 vs SUV39H1Δ/H2Δ clones, represented in 1 kb bins. The red traces represent the control samples Neo4p13 EP (solid red) and SUV39H1Δ/H2Δ EP (light red). The grey trace represent the combined mean enrichment for three groups of Day 100 clones: BR1, BR2, and BR3, with Day 100 +/− 1 standard deviation (SD, light grey) **(F)** CENP-A enrichment in three genomic regions surrounding the Neo4p13 locus: the initial Neo4p13 region (42.868–42.958 Mb) and flanking outside regions 90 Kb upstream (“Left (−90 Kb)”) and downstream (“Right (+90 Kb)”). Left, Enrichment values are normalized to the Neo4p13 early passage (EP) control sample (baseline = 1). Whiskers represent the standard deviation (SD), with colours highlighting the genomic regions and baseline normalization (dashed line at 1) are included for clarity. Groups of 100-day biological replicates (BR1, BR2, BR3) are compared. Right, Relative CENP-A enrichment as in left panel but normalized to the SUV39 H1/H2Δ early passage (EP) control sample. **(G)** CENP-A CUT&RUN of early passage parental Neo4p13 (red) versus SUV39H1Δ/H2Δ/SUZ12 Δ (red) EP clones (red) versus corresponding 100-day clones (grey). compared with panel D above. **(H)** Line graph as in E but comparing WT Neo4p13 vs SUV39H1Δ/H2Δ/Suz12Δ clones. **(I)** Left: Relative CENP-A enrichment analogous to panel F, normalized to the WT Neo4p13 EP. Right: Relative CENP-A enrichment normalized to SUV39H1Δ/H2Δ/SUZ12Δ EP.

To assess their contributions, we targeted lentiviral Cas9 to the genes encoding SETDB1, SUV39H1 and SUV39H2 to make SETDB1 null (SETDB1Δ) and SUV39H1/H2 double null mutant lines (SUV39H1Δ/H2Δ) (Fig 3B). Loss of SUV39H1/H2 resulted in a modest reduction in global H3K9me3 levels, likely due to other H3K9 methyltransferases (Fig 3B). In contrast, SETDB1 loss did not affect global H3K9me3 levels detectable by immunoblot (Fig 3B). Using H3K9me3-directed CUT&RUN, we found that deletion of SUV39H1/H2 but not SETDB1, resulted in the near-complete loss of H3K9me3 from the neocentromere domain (Fig 3C). This depletion resulted in a substantial ∼20 kilobase drift of CENP-A positioning in a telomere-proximal direction following the generation of the mutant cells, clonal isolation, and expansion (∼50 days in culture) (Fig 3D-F). To assess long-term adaptation, we employed the clonal culture strategy as outlined in Fig 1. We find that, following an additional 100 days of continuous proliferation, SUV39H1Δ/H2Δ clones had all moved to a more telomere-proximal position compared to wild-type Neo4p13 (Fig 3D-F). Moreover, CENP-A enrichment is generally higher across all clones versus Neo4p13 EP (Fig 3D-F). Importantly, despite an initial expansion in early passage clones, the overall size of the CENP-A domain remained largely consistent at 90-110 kb in evolved clones, suggesting another level of size control that maintains neocentromere size, preventing spreading beyond a confined locus.

Polycomb Repressive Complex 2 is known to compensate for H3K9me3 loss in some SUV39 mutant contexts^48^. Indeed, we observe an increase in PRC2-mediated H3K27me3 facultative heterochromatin compensating for loss SUV39-mediated loss of H3K9me3 at 4p13 (Fig 3C), as previously observed on H3K9me3-depleted HACs^49^. To remove H3K27me3, we disrupted the gene encoding Suppressor of Zeste 12 (SUZ12), an essential component of the PRC2 complex (Fig 3B)^50^. Deletion of SUZ12 resulted in a complete loss of H3K27me3 in SUV39H1Δ/H2Δ/SUZ12Δ triple mutant Neo4p13 cells (Fig 3B, C). Like SUV39H1Δ/H2Δ, removal of H3K27me3 resulted in a substantial enrichment of CENP-A compared to the control (Fig 3G, H and I). Strikingly, size control is largely lost in this context, resulting in the CENP-A domain doubling in size compared to the original Neo4p13 locus (Fig 3G, H and I), moving much further than SUV39H1Δ/H2Δ to a size up to 150 kilobases and breaching into the nearest gene boundary (GRXCR1). Combined, these results indicate that heterochromatin marked by H3K9me3 and H3K27me3 can form clear functional boundaries at this neocentromere, and maintain tight size control of the CENP-A domain, independent of 5mCG methylation and alpha satellites.

### Canonical alpha satellite-based centromeres are characterized by heterochromatin dips

Our analysis of a neocentromere highlighted the importance of heterochromatin in forming CENP-A boundaries. Next, we investigated whether similar boundary mechanisms exist at canonical centromeres or if canonical centromeres are controlled differently. Characteristic features of canonical centromeres, such as repetitive alpha satellite DNA and high levels of DNA methylation, may also contribute to maintaining CENP-A positioning. However, the characterisation of centromeric chromatin organisation at canonical centromeres has thus far been limited due to the highly repetitive nature of satellite DNA, preventing the reliable mapping of short-read sequencing such as CUT&RUN. To circumvent this issue, we employed ONT ultra-long-read DiMelo-seq in combination with the isogenomic RPE reference^39^ to accurately map CENP-A and H3K9me3 occupancy at canonical centromeres (Fig 4A)^38^. We performed this analysis in the same RPE1-Neo4p13-bearing cell line for consistency across our experiments. As ONT enables direct genomic DNA sequencing lacking PCR amplification, we can determine the single-molecule occupancy of our target of interest across centromeres/pericentromeres.

**Figure 4:**
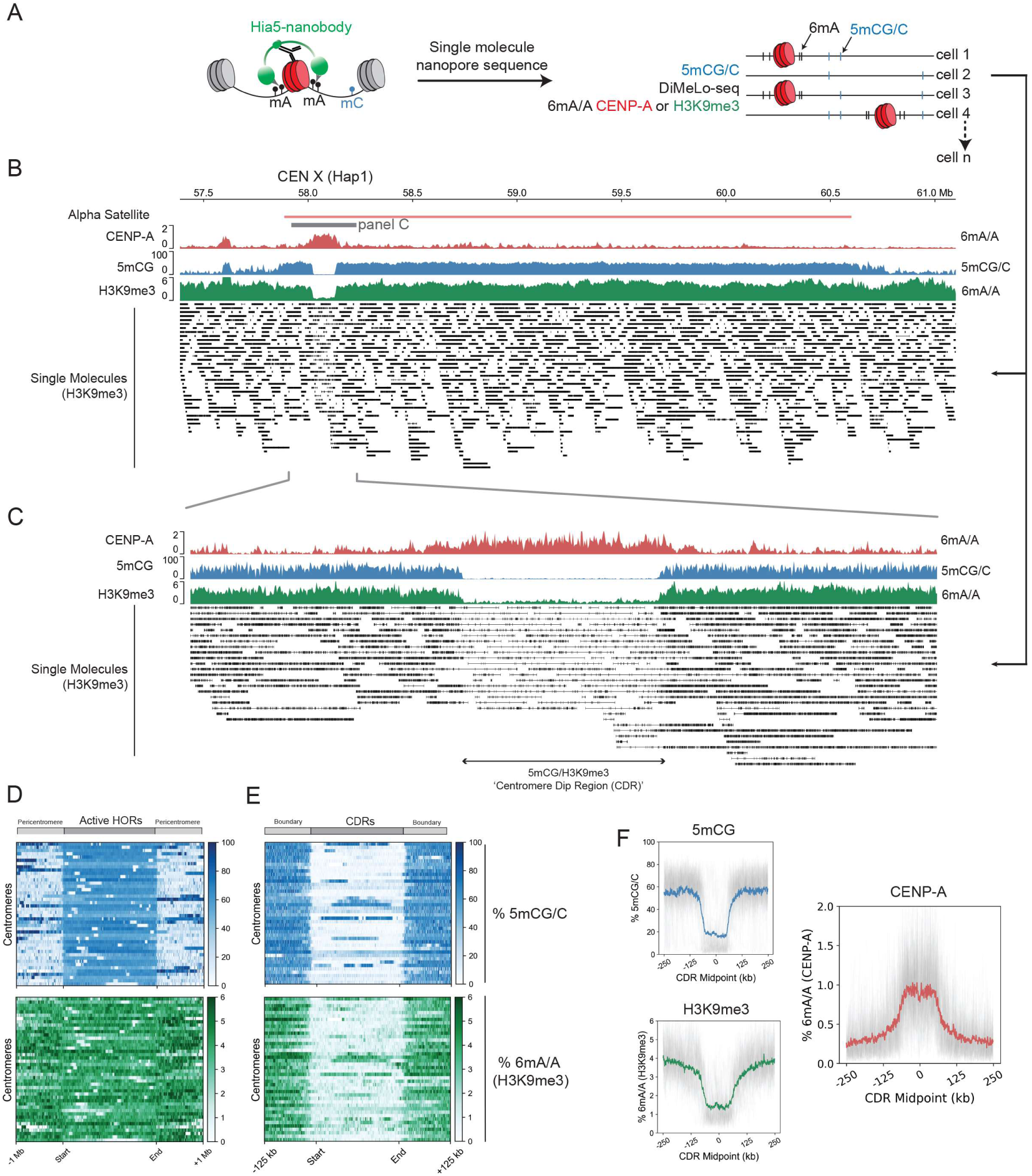
Heterochromatin dips are characteristic of canonical centromeres. **(A)** Diagram of DiMeLo-Seq workflow with averaged 6mA/A probabilities and single molecule alignments. Vertical lines of each read represent CENP-A or H3K9me3-directed exogenous 6mA. **(B)** View of the entire HOR of CEN X (Hap 1), with DiMelo-Seq enrichments for CENP-A (red) and H3K9me3 (green), as well as 5mCG enrichment (blue). Single molecules of each long read for H3K9me3 (black). Specifically where CENP-A resides, there is the characteristic dip in DNA methylation (5mCG), but also H3K9me3 heterochromatin. **(C)** A closer view of CEN X, showing the dip in H3K9me3 and boundaries surrounding the active CENP-A-containing region. Single molecules at this site contain largely unmethylated adenine-called reads. **(D)** Heatmaps illustrating the distribution of 5mCG (top, blue) and H3K9me3-directed 6mA (bottom, green) modifications across centromeric Active HOR regions. Each line on the heatmap spans the Active HOR of a single centromere and flanking regions ±1 Mb upstream and downstream. All Active HOR regions have been scaled to the same length to allow direct comparison of methylation patterns across all centromeres. Signal intensity is represented as the mean methylated fraction across fixed genomic bins, with darker shades indicating higher modification levels. Start and end of the HOR regions are indicated, highlighting methylation patterns in and around these active centromeric domains. **(E)** Heatmaps as in D but specifically across Centromere Dip Regions (CDRs), as defined by the boundary of 5mCG dips on either side of each CENP-A-containing region. All CDRs have been scaled to the same length. **(F)** Individual (grey) and aggregated (coloured) modification traces of indicated signals across all CDRs, centred on CDR midpoints, with ±250 Kb flanking regions.

First, to determine the specificity of the CENP-A 6mA signal, we utilised an endogenous homozygous knock-in AID-eYFP-CENP-A hTERT-RPE1 cell line to auxin deplete all endogenous CENP-A^51^. We carried out a 3-hour IAA treatment followed by light fixation and CENP-A DiMeLo-seq (Fig S2A). CENP-A was effectively depleted from cells (Fig S2B) and the corresponding CENP-A 6mA signal was specifically eliminated from all CDRs, indicating 6mA is CENP-A specific and providing us with a true background CENP-A 6mA/A signal (Fig S2C, E and F). Consistent with previous observations^3,29,39^, active alpha satellite Higher Order Repeat (HOR) arrays generally contain high levels of DNA methylation (Fig 4B-E, blue). Moreover, active centromeric regions containing CENP-A feature the previously characterised Centromere Dip Regions (CDRs), depleted of 5mCG, which correlate directly with all CENP-A-containing regions (Fig 4B, C, E and F). Interestingly, 5mCG levels were unaffected by acute loss of CENP-A (Fig S2D), indicating that the local dip in DNA methylation, is relatively stable, at least within a few hours of CENP-A depletion.

H3K9me3 is normally present in high densities in pericentric regions in classical constitutive HP1-mediated heterochromatin domains^52–54^. To determine the resolution of our H3K9me3 DiMeLo-seq, we first assessed H3K9me3 in pericentric regions, where H3K9me3 levels are normally high, specifically using HSat3 as an example (Fig S3A). We can achieve near-single-nucleosome resolution for H3K9me3, giving us an estimate of methylated nucleosome densities in these regions that is consistent with the reported average chromatin repeat unit length for HSat3 (Fig S3B and C)^55^.

In the classic centric/pericentric dichotomy, heterochromatin, marked by H3K9me3, is considered to be largely pericentric, whereas the kinetochore-forming CENP-A domain is largely depleted of heterochromatin^31,32,36,56^. Moreover, centromeres on human artificial chromosomes resist silencing mediated by H3K3me3/H3K27me3^49^, and H3K9me3 is not required for CENP-A-driven epigenetic memory ^57^, suggesting an inverse relationship and functional boundary potential. However using DiMeLo-seq, it has recently been unambiguously shown that H3K9me3 is also present within HORs that form a template for CENP-A^38^. Consistent with these data, we find high levels of H3K9me3 across all pericentromeres and active HORs in RPE1 (Fig 4B, C and D, green). Importantly, we identify a distinct depletion in H3K9me3 where CENP-A resides at all centromeres, forming a tight boundary around the CENP-A domain in a manner analogous to 5mCG (Fig 4C, E and F and Fig S4). Thus, this dataset identifies the existence of a heterochromatin dip region (HDR) overlapping with the previously identified CDR defined by a lack of DNA methylation.

### Centromeric H3K9me3 is differentially controlled by SETDB1, SUV39, and non-canonical SUZ12 functions

To uncover the significance of the H3K9me3 boundaries at canonical alpha satellite-based centromeres, we sought to manipulate these and determine whether centromere size control and migration are affected. To this end, we assessed H3K9me3 levels at canonical centromeres in the SETDB1Δ, SUV39H1Δ/H2Δ and SUV39H1Δ/H2Δ/SUZ12Δ double and triple mutants described above. We find that in SUV39H1Δ/H2Δ double mutants, H3K9me3 levels are decreased across both the flanking pericentromere and within centromeric active HORs, compared to parental cells (Fig 5A, C). Surprisingly, we find that SETDB1 contributes to H3K9me3 deposition specifically within the centromeric HORs and not in the flanking pericentric regions (Fig 5A, C), indicating it has centromere-specific roles distinct from SUV39 enzymes. Furthermore, we find that SUV39H1Δ/H2Δ/SUZ12Δ triple mutants offered the most prominent effect on H3K9me3. Whilst levels were reduced in the pericentromeric regions to similar levels as in SUV39H1Δ/H2Δ double mutant cells, there was a striking and substantial loss of H3K9me3 that specifically affected active HORs, even beyond that observed in SETDB1Δ cells (Fig 5A, C). Hence, whilst SUV39H1/H2Δ generally reduces H3K9me3 in peri and alpha-satellite containing chromatin, SETDB1 and SUZ12 deletion appear to have selective effects on alpha-satellite H3K9me3.

**Figure 5:**
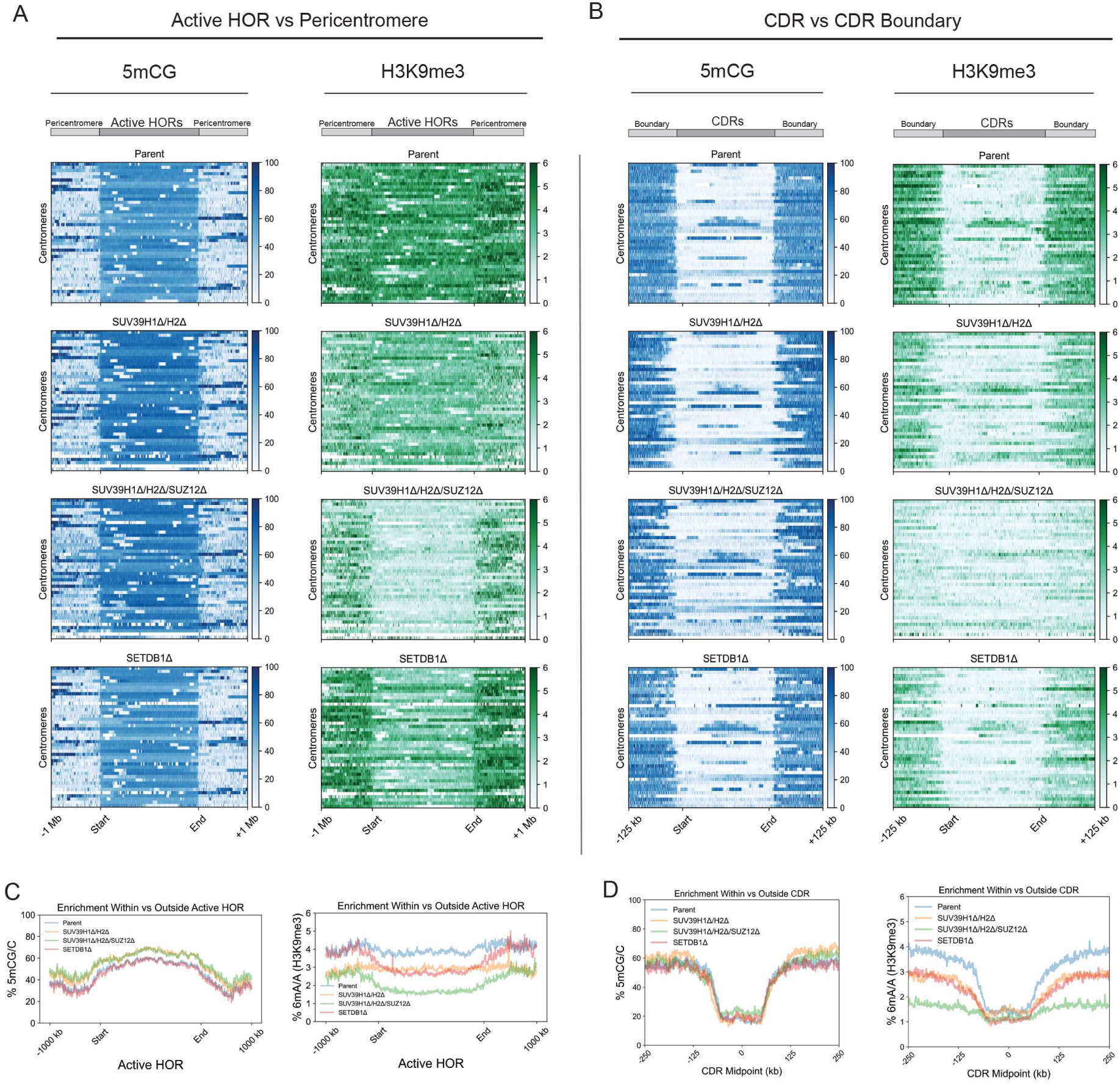
Centromeric H3K9me3 is regulated by a combination of SETDB1, SUV39 and non-canonical SUZ12 functions. **(A)** Heatmaps for WT, SUV39H1Δ/H2Δ, SUV39H1Δ/H2Δ/SUZ12Δ and SETDBΔ cell lines, illustrating the distribution of 5mCG (blue) and H3K9me3-directed 6mA (green) modifications across centromeric Active HORs. Each line on the heatmap spans the Active HORs of a single centromere and flanking regions ±1 Mb upstream and downstream. All Active HOR regions have been scaled to the same length to allow direct comparison of methylation patterns across all centromeres. **(B)** Heatmaps as in B but specifically across Centromere Dip Regions (CDRs), as defined by the boundary of 5mCG dips on either side of each CENP-A-containing region. All CDRs have been scaled to the same length. **(C)** Average indicated modification signal across all Active HOR regions (scaled equally), flanked by ±1000 Kb. **(D)** Average indicated modification signal across all CDRs regions flanked by ±250 Kb.

We then analysed the CENP-A-containing DNA methylation-depleted regions (CDRs). Here, we find local H3K9me3 boundaries to be effectively eliminated in SUV39H1Δ/H2Δ/SUZ12Δ backgrounds, with visible reductions in boundary strength also apparent in SUV39H1Δ/H2Δ and SETDB1Δ respectively (Fig 5B, D). While these mutants substantially lose H3K9me3, the relative levels of 5mCG across active HORs were higher in SUV39Δ contexts and unaffected in SETDB1Δ background. The loss of these enzymes had only a minor effect on the 5mCG-defined CDR boundaries (Fig 5B, D). Therefore, in this context, we are primarily assessing the contributions of H3K9me3 to centromere position, largely independent of the role of DNA methylation.

### Heterochromatin is the primary functional CENP-A boundary at canonical centromeres

Next, having established means to selectively deplete heterochromatin from active HORs and pericentric HSats, we assessed its contribution to CENP-A positioning. In parental RPE-Neo4p13 cells, we observed a tight positioning of CENP-A within the CDR, with levels rapidly dropping off outside of the 5mCG CDR boundaries (Fig 4F, Fig 6A, B and D). In both SETDB1Δ and SUV39H1Δ/H2Δ/SUZ12Δ backgrounds, we observe a significant overall expansion of CENP-A following ∼50 days in culture (time to generate these knockouts) (Fig 6A, B an F). The majority of centromeres had moved beyond their original CDR boundaries of parental cells – most prominently in the SUV39H1Δ/H2Δ/SUZ12Δ triple mutant cells where H3K9me3 levels are the most reduced. The CENP-A enrichment ratio within vs outside the CDR in control samples drops in both mutants (Fig 6D), even below a ratio of 1 in many centromeres, indicating that CENP-A has become more enriched in the surrounding 100 kb region outside of the originally defined centromere boundaries than within. Interestingly, the DNA methylation dips were largely maintained, even if H3K9me3 levels were uniformly low across the CDR and adjacent HORs (Fig 5C, 6A, E and F). Nevertheless, we could observe erosion of 5mCG at the centromere boundaries indicating that H3K9me3 and H3K27me3 methyltransferases have an indirect effect on DNA methylation (Fig 6E). In summary, the resulting expansion of CENP-A correlates with the loss of H3K9me3 and, to a lesser extent, with reduction of DNA methylation at the boundary. These observations indicate that H3K9me3 heterochromatin is the primary functional boundary maintaining CENP-A position.

**Figure 6:**
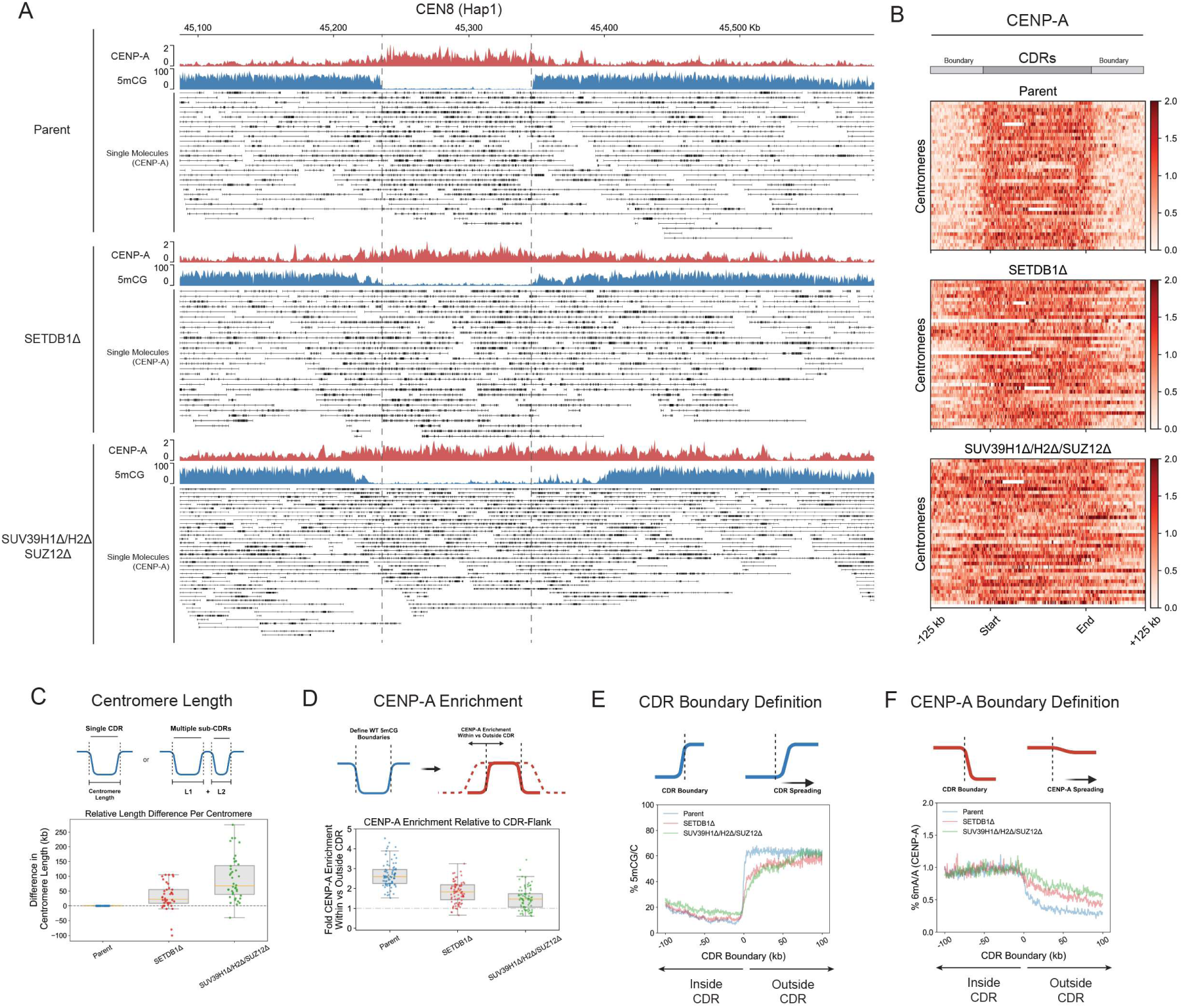
Heterochromatin forms a functional boundary at canonical centromeres. **(A)** View of the entire HOR of CEN 8 (Hap 1) for parental (RPE-Neo4p13), SETDB1Δ and SUV39H1Δ/H2Δ/SUZ12Δ with aggregate DiMelo-Seq enrichments for CENP-A (red) and 5mCG enrichment (blue). Single molecules alignments for CENP-A-directed 6mA (black). **(B)** Heatmaps for each mutant illustrating the distribution of CENP-A (red) across Centromere Dip Regions (CDRs), as defined by the boundary of 5mCG dips on either side of each CENP-A-containing region in the parent cell line (RPE-Neo4p13). Each line on the heatmap represents a single CDR/Centromere. All CDRs have been sorted numerically and scaled to the same length for comparison. **(C)** Centromere length as defined by the difference in sum total length of CDRs/sub-CDRs relative to the corresponding parental (RPE-Neo4p13) centromere. Box plots summarize the distribution for parent and mutant lines, while individual data points are shown as coloured dots. A dashed reference line at 1 indicates parental centromere length. The orange line represents the median centromere length. **(D)** CENP-A Enrichment: Box plot depicting the ratio of CENP-A enrichment within CDRs relative to their flanking regions (±100 Kb) for parent and mutant lines. Individual data points are shown as coloured dots, while box plots represent the distribution of ratios. A dashed reference line at 1 indicates equal enrichment within and outside the HORs. The orange line represents the medium CENP-A ratio. **(E)** CDR boundary definition: plot of all 5mCG CDR boundary profiles centred on CDR boundaries (±100 Kb). Mean of combined aggregated traces is shown for all samples. **(F)** CENP-A boundary definition: Plot of CENP-A enrichment inside versus outside the CDR boundary (defined by WT 5mCG) for each cell line (±100 Kb). Mean of combined CENP-A (6mA/A) aggregated trace is shown for all samples.

### CENP-A drift at canonical centromeres over time is exacerbated by heterochromatin defects

Given our observations that Neo4p13 drifts gradually over time (Fig 1D), we sought to determine whether canonical centromeres display a similar behaviour. To test this, we picked a clone in which the CENP-A domain at the Neo4p13 locus has shifted significantly following 100 days of proliferation (clone 1.2; Fig 1D), and compared global centromere positioning to the early passage parental line. Similarly, we picked a SUV39H1Δ/H2Δ/SUZ12Δ clone in which the 4p13 neocentromere also moved significantly over a 100-day culture period (clone 1.2; Fig 3G). We then performed CENP-A DiMelo-seq to determine any differences in CENP-A position and size over time. Similar to the neocentromere, canonical centromeres move subtly with time over the 100-day period (Fig S5). Parental centromere length (Fig S5C) and CDR boundary definition (Fig S5E) are largely maintained or subtly eroded after 100 days. CENP-A position begins to gradually shift from the initial centromere boundary (Fig S5D and F). In contrast, already in early passage SUV39H1Δ/H2Δ/SUZ12Δ cells as well as in 100-day clones, the CENP-A domain has shifted significantly beyond any drift observed in parental cells, even following 100 days of proliferation (Fig S5D and F). In addition, centromere length has expanded (Fig S5C), and CDR definition has significantly eroded (Fig S5E) in these clones. We also observe cases where CENP-A position has almost entirely vacated the parental position in the SUV39H1Δ/H2Δ/SUZ12Δ 100-day background (Fig S5A and B). Interestingly, here 5mCG methylation begins to reoccupy the initial CDR in the absence of CENP-A (Fig S5A, bottom). This suggests that in addition to our observation at the neocentromere, where seeding CENP-A coincides with a loss of DNA methylation, the inverse can also happen where the gradual loss of CENP-A results in the re-establishment of DNA methylation. Moreover, our assessment of temporal CENP-A domain dynamics, indicates that the dramatic movement of CENP-A in methyltransferase mutants is the result of the loss of these enzymes and not simply a consequence of clonal selection and long-term culture.

### Loss of heterochromatin within HORs results in new CDRs and ectopic functional CENP-A domains

In addition to our assessment of the immediate CENP-A boundaries, we further analysed the contribution of heterochromatin to CENP-A occupancy across the entire active HOR of each centromere. Surprisingly, we observed many cases where the loss of heterochromatin resulted in new CENP-A peaks distal to the primary CENP-A domain, but within the same alpha satellite HORs. These peaks correlate with loss of 5mCG and H3K9me3 thus forming characteristically similar chromatin features to the primary active centromere/CDR. For example, CEN1 Hap1 and CEN9 Hap1 of SUV39H1Δ/H2Δ/SUZ12Δ both contain new CDRs ∼3 Mb away from the original primary CDR (Fig 7A). SETDB1Δ cells similarly show specific examples of new CDRs on CEN20 Hap1 and CEN19 Hap2 (Fig 7A). To thoroughly assess all new CDRs, we used CDR-Finder^58^ followed by manual curation to ensure all CDRs are CENP-A containing (Table 1, see methods for details). Indeed, SUV39H1Δ/H2Δ/SUZ12Δ and SETDB1Δ have numerous new CENP-A containing CDRs which have formed distal on the same alpha satellite HOR (Table 1). Furthermore, a new CENP-A peak is also present on CEN21 Hap1 of parental cells following 100-day culture, suggesting new CENP-A domain seeding could be a general phenomenon that occurs at low frequency even in wild-type cells over time. Some HOR regions appear to be preferred sites for new CENP-A domain formation across our cell lines (e.g. Chrs 9, 20 and 21). No new CDRs appeared in SUV39H1Δ/H2Δ cells.

**Figure 7:**
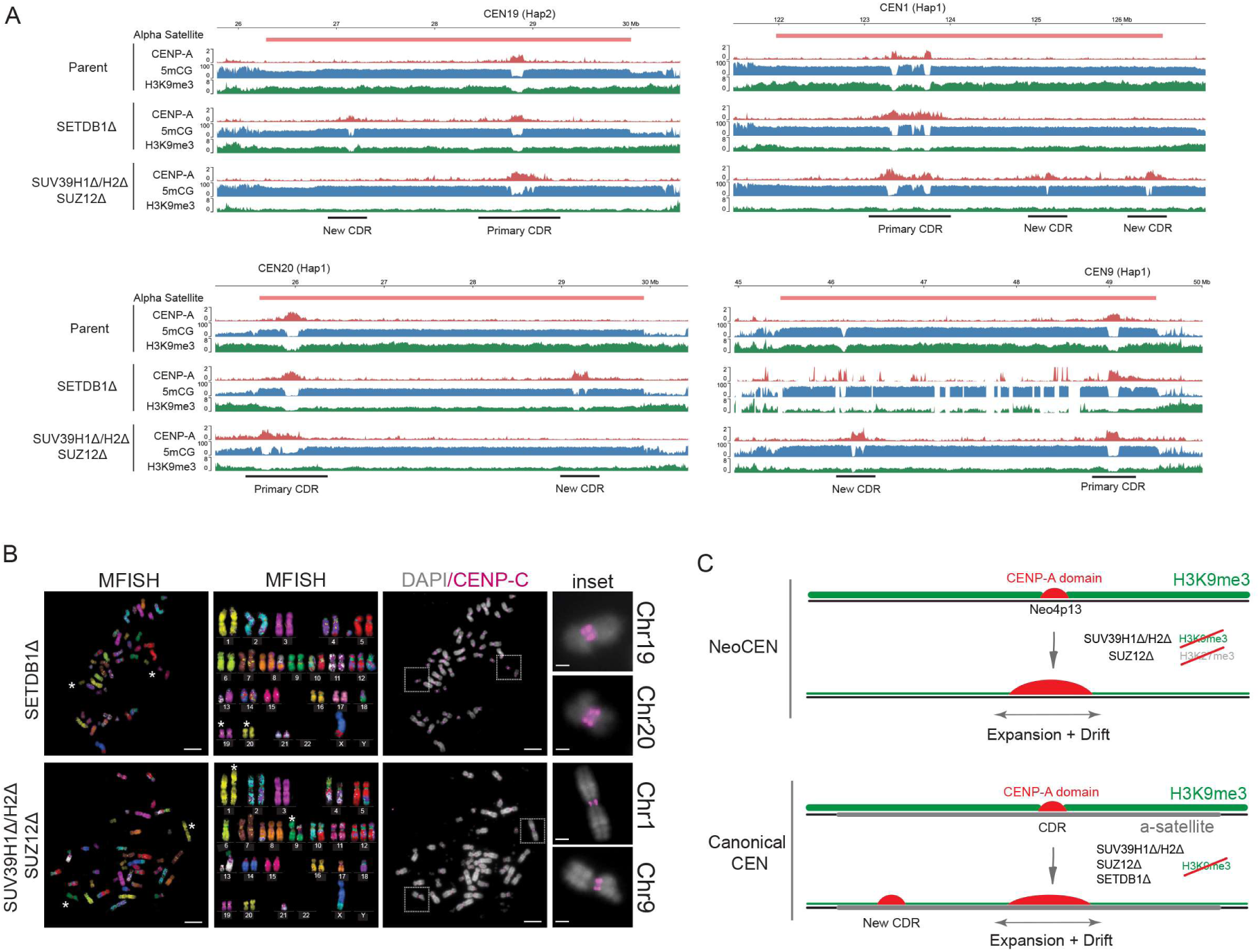
Nucleation of new CENP-A peaks and CDRs, forming functional dicentrics. **(A)** Representative genome tracks for the entire alpha-satellites of CEN 1 (Hap1), CEN 9 (Hap1), CEN20 (Hap1) and CEN 19 (Hap2), showing CENP-A (red), 5mCG (blue) and H3K9me3 (green) for Parental, SUV39H1Δ/H2Δ/SUZ12Δ and SETDBΔ cell lines. The primary and newly-formed centromeres are highlighted in each case, corresponding with new CENP-A seeding and 5mCG and H3K9me3 dips. **(B)** Representative IF-mFISH images of metaphase spreads, showing karyotype and dicentric chromosomes for SETDB1Δ and SUV39H1Δ/H2Δ/SUZ12Δ cells as per (A) above, identified by CENP-C (magenta), and chromosome-specific mFISH paint. Scale bar 10 μm; Inset scale bar 5 μm **(C)** Model summarising the differential role of H3K9me3 methyltransferases at neocentromeres and canonical centromeres and their role in defining CENP-A domain size, position and number. Left, Loss of heterochromatin (H3K9me3/H3K27me3) at Neo4p13 results in expansion and drift of the neocentromere. Right, Loss of H3K9me3 at canonical alpha satellite centromeres leads to centromere expansion, drift and nucleation of new centromeres at distal sites on the alpha satellite.

We find all mutant lines to generally maintain a diploid karyotype as measured by DNA content (Fig S6). Using spectral karyotyping we identify some translocations and aneuploidies of different chromosomes across each cell line (Fig S7A and B). However, loss of heterochromatin may have pleiotropic effects and, generally, we do not find that the translocations result from centromeric breakages and appear to occur independently from new CENP-A peaks. We do observe some instances of HOR destabilisation not observed in parental cells. An example is CEN9 Hap 1 in SETDB1Δ (Fig 7A). However, the majority of HORs appear stable and translocation events appear not to be directly linked to centromere defects.

Finally, to determine whether the ectopic CENP-A peaks form distinct centromere domains, we sought to directly visualize these new centromeres on mitotic chromosomes. To enable resolving distal CENP-A domains, we chose centromeres where the distance between CENP-A peaks was greater than 2 Mb. Using a combination of multicolour FISH (mFISH) and immunofluorescence for CENP-C, we can resolve dicentric chromosomes in metaphase spreads for chromosomes 1 and 9 of SUV39H1Δ/H2Δ/SUZ12Δ mutants as well as chromosomes 19 and 20 in SETDB1Δ cells. Here, CENP-C appears as distinct double loci at these centromeres (Fig 7B), corresponding to the new DiMeLo-seq peaks (Fig 7A), indicating that these loci form functional dicentrics with two centromeric sites within the same alpha satellite array.

## Discussion

### Heterochromatin forms a centromere boundary that controls centromere position and size

Constitutive heterochromatin (marked by H3K9me3) flanking the centromeric HORs, offers a potential boundary to maintain the overall centromere position^30^. Furthermore, the recent development of DiMeLo-seq has shown that H3K9me3 spans both pericentric and centromeric alpha satellites^38^. This poses questions as to 1) How H3K9me3 is regulated at alpha satellites, and 2) whether it is functionally involved in maintaining centromere position/boundaries.

We dissected the role of H3K9me3 at the centromere and pericentromere, both at a unique neocentromere as well as canonical centromeres. We discovered that human neocentromere Neo4p13^18^ is enriched in SUV39H1 and SUV39H2-dependent H3K9me3 but is formed in generally low levels of DNA methylation and lacks alpha satellites, indicating that the latter are not essential contributors to centromere positioning, at least at this neocentromere. Further, seeding of CENP-A induces local depletion of heterochromatin, specific to the haplotype with the neocentromere and precisely surrounding the 90 kb CENP-A domain. We previously estimated around 100 CENP-A nucleosomes per haplotype in RPE1 cells^22^. Within a ∼100 kb centromere domain, this translates in 1/5 nucleosomes to be CENP-A containing. Thus, while the loss of H3K9me3 is explained in part by the displacement of histone H3 by CENP-A, our findings suggest that CENP-A seeding also results in the active removal of H3K9 trimethylation. We find that experimental removal of heterochromatin (H3K9me3 and compensating H3K27me3) within this domain results in a drift of the CENP-A domain and significantly expands its size. We hypothesized that these heterochromatin marks are maintained through self-templated propagation forming a competitive, incompatible barrier for CENP-A deposition.

We find that local depletion of H3K9me3 chromatin extends to canonical centromeres where the CENP-A domain not only correlates with a dip in CpG methylation but also a corresponding dip in H3K9 trimethylation. Again, perturbation of these H3K9me3 boundaries results in a substantial movement and expansion of CENP-A domains at all centromeres within several cell divisions. Importantly, while loss of H3K9me results in some erosion of CpG methylation at the CDR boundary, the DNA methylation dips are largely maintained. Altogether these results indicate that DNA methylation may not act as the primary and sole boundary restricting the CENP-A domain but that instead, heterochromatin is key to restricting the functional CENP-A domain, at least at the long timescales we have analysed. However, we cannot exclude that rapid and/or complete perturbation of the DNA methylation flanking the CDR could affect CENP-A position, possibly even without affecting the surrounding heterochromatin, as loss of DNA methylation may have only a minor impact on the heterochromatin level^59^. Most likely, while H3K9me3 appears dominant in the depletion setting presented here, DNA methylation and H3K9 methylation ultimately act in tandem at the centromere. Determining their individual roles requires future separation-of-function studies.

### Differential regulation of centromeric H3K9me3

Centromeric and pericentromeric repeats are largely repressed by H3K9me3^38,52,53^. The classical model for the regulation of pericentric heterochromatin involved the deposition of H3K9 methylation by the SUV39 methyltransferases coupled to Heterochromatin Protein 1 (HP1)-mediated repression^52^. How H3K9me3 is regulated within the centromere has remained unclear. Our data supports a general role for SUV39 methyltransferases at both pericentric HSat sequences as well as centromeric alpha satellites. Further, we discovered that the SETDB1 acts as a methyltransferase specific to active alpha satellite HORs. SETDB1 is known to silence transposable elements^45,46 47^. Interestingly, active HORs, like TEs are highly methylated compared to adjacent inactive arrays^3,29^. Possibly, the high degree of methylation contributes to SETDB1 recruitment. Furthermore, at TEs, SETDB1 is recruited via transposon-encoded sequence elements^45,46^. Understanding whether cryptic sequences within active HORs contribute towards SETDB1 targeting is an important future goal.

Surprisingly, we also identified a significant role for the PRC2 component SUZ12 in the regulation of centromeric satellites, with a substantial further depletion of H3K9me3 at active HORs upon loss of SUZ12. Whilst the interplay between H3K9me3 and H3K27me3 is relatively poorly understood, it has previously been shown that SUZ12 regulates histone H3 lysine 9 methylation and HP1 alpha distribution in flies^60^. Furthermore, SETDB1 and SUZ12 have been found to frequently colocalize in mouse ES cells^61^, as well as SUV39^60^. While we cannot exclude secondary effects of long-term gene deletions in this study, these observations suggest a possible direct functional interaction between SETDB1, SUV39 and SUZ12 at centromeric repeats which requires further careful biochemical and genetic dissection.

### Heterochromatin limits the number of CENP-A/CDR domains on alpha satellite HORs

Why do centromeric satellites feature such a high level of repressive chromatin surrounding the active centromeric locus? One reason might be to retain a tight concentration of CENP-A at a defined locus. Prior evidence suggests that centromere nucleation is a concentration-dependent mechanism^5,6^, requiring a critical concentration of centromere proteins to trigger an active centromere state. The same principle holds true for kinetochore formation^62^. Indeed, CENP-A is loaded onto chromatin at low concentrations outside the centromere but removed through DNA replication apart from at the centromere^22,63,64^. Therefore, if the critical CENP-A concentration is not retained within a small, defined region between relatively strict boundaries, the centromere may spread, even within the megabase-arrays of alpha satellite DNA, and may not reach its critical concentration at the primary site with the same efficiency. Our discovery of new CENP-A domains distal to the primary CDR, that appear to form functional centromeres in heterochromatin-deficient mutants, is consistent with a role for heterochromatin (and DNA methylation), not only in acting as a boundary at existing CENP-A domains but also to suppress aberrant centromere formation at regions outside of the primary active centromere locus. One possible role of H3K9me3 and DNA methylation may be to constrain centromeric transcription specifically within CDR where CENP-A assembles. Given that centromeric transcription appears to be required for CENP-A function^65^, and is indeed activated upon centromere repositioning^66^; it is tempting to speculate that transcripts or indeed RNA polymerase II are confined to these CDRs by repressive chromatin.

The new CENP-A domains we observe in heterochromatin mutants occur within the HOR and are thus generally positioned relatively close to the primary CENP-A domain. Hence, the 1-3 Mb distances that separate the primary and newly formed CDRs appear to act as functional dicentrics that are generally retained and permissive during mitosis. This is consistent with previous work that showed that two active centromeres, even when several tens of megabases apart can be mitotically stable^67^.

We cannot exclude the possibility that more distal CENP-A domains appear in these mutants but are lost due to selection. Furthermore, it is possible that the new CDRs form at some frequency on HORs even in wild type cells, but are tolerated due to the large size of the alpha satellite array. Centromeres from different individuals can be at drastically different genomic loci on their respective HORs^3,68^. Therefore, centromere positioning within the HOR may be to some extent functionally flexible and contribute to the overall robustness of centromere function and evolution. In sum, we identified heterochromatin as an important contributor to maintaining CENP-A boundaries. How these boundaries are set at the remarkably constant size of about 100 kilobases, independent of underlying sequence context, remains an open question. Possibly, the assembly of the multiprotein centromere complex and the functional kinetochore ties into control mechanisms dynamically dictating the number of CENP-A nucleosomes. Understanding the dynamic feedback between CENP-A chromatin, its boundaries, and the centromere/kinetochore complex is an important future goal.

## Materials and Methods

### Cell line and Culture Conditions

All human cell lines were grown at 37 °C, 5% CO2. RPE1 cell lines were grown in DMEM/F-12 media (Gibco) supplemented with 10% fetal bovine serum (FBS) (Gibco), 1x non-essential amino acids (NEAAs) (Gibco), 2 mM glutamine (Gibco), 100 U/ml penicillin (Gibco) and 100 µg/ml streptomycin (Gibco). HEK293 cells were grown in DMEM (Gibco) supplemented with 10% FBS (Gibco).

### Cell Line Generation and Lentiviral Transduction

The RPE-Neo4p13 cell line was generated as previously described^18^. Deletion of genes encoding SUV39H1/H2 and SETDB1 in the RPE-Neo4p13 cell line was performed by transfection of LentiCRISPRv2-Blast (Addgene #83480). gRNAs were designed (Table 1, below) and cloned according to standard lentiCRISPRv2 cloning protocol^69^. For lentiviral generation, HEK293T cells were seeded at a density of 1 × 10⁶ cells in DMEM supplemented with 10% FBS in a T25 flask one day prior to transfection. Cells were gently refreshed with serum-containing, antibiotic-free medium prior to transfection. A plasmid mixture (0.62 µg pMD2 (Addgene #12259), 1.87 µg psPAX2 (Addgene #12260), and 2.5 µg pLentiCRSPRv2) was diluted in 100 µL Opti-MEM, and 10 µL Lipofectamine LTX was diluted in another 100 µL Opti-MEM. The DNA and Lipofectamine solutions were combined, mixed gently, and incubated for 5 minutes at room temperature before being added dropwise to the T-25 flask. After 24 hours at 37°C, the medium was replaced, and cells were cultured for an additional 48 hours. The conditioned medium was then harvested, filtered through a 0.45 µm filter, and either frozen at −80 °C or used immediately to infect RPE-Neo4p13 cells. For transduction, RPE-Neo4p13 cells were seeded at 5 × 10^4^ cells per well in a 6-well plate the day before infection. Filtered viral supernatant was mixed with antibiotic-free medium, supplemented with 3.2 µL of 10 mg/mL Polybrene, and added to the cells for 48 hrs. After infection, cells were washed four times with PBS, replaced with regular medium, and, if required, treated with the 5 µg/ml Blasticidin for 5 days. Cells were then clonally sorted into 96-well plates, grown out and screened by immunoblot for positive clonal knockouts for each target. For generation of SUV39H1Δ/H2Δ/SUZ12Δ cell line, SUZ12 gRNAs (Table 1, below) were designed and cloned into pSpCas9(BB)-2A-GFP (PX458) plasmid using the standard Zhang lab gRNA insertion protocol^70^. Clonal SUV39H1Δ/H2Δ cells were transiently reverse co-transfected with both PX458 using Mirus® TransIT-LT1 protocol at a 2:1 Reagent to DNA ratio in a 6-well plate at a density of 6 × 10^4^ cells per well. After 48 hrs, cells were sorted for high GFP expressing into a polyclonal population and allowed to recover for one week. When cells were proliferating, cells were clonally sorted (GFP negative) into 96-well plates and allowed to expand, followed by screening by immunoblot for clones in which both SUZ12 and resulting H3K27me3 were lost.

### FACS/Flow Cytometry and Cell Cycle Analysis

For fluorescence-activated cell sorting (FACS) and cytometry, cells were harvested by centrifuging at 500 x g for 5 mins and re-suspended in ice-cold sorting medium, which consisted of 1% fetal bovine serum in PBS, supplemented with 0.25 mg/mL Fungizone (Thermo Fisher Scientific) and 0.25 μg/mL Amphotericin B and 10 μg/mL Gentamicin (Gibco). The cell suspension was filtered through 5 mL polystyrene round-bottom tubes equipped with cell-strainer caps (Falcon) prior to sorting and analysis using the FACSAria III or FACSAria Fusion Cell Sorter (BD Biosciences). Sorted cells were collected into 96-well plates containing conditional medium, prepared as a 1:1 mix of fresh complete medium and 0.45 μm-filtered medium from proliferating cell cultures, supplemented with 20% fetal bovine serum, 0.25 mg/mL Fungizone (Thermo Fisher Scientific), and 0.25 μg/mL Amphotericin B and 10 μg/mL Gentamicin (GIBCO).

For cell cycle analysis, cells were harvested and washed with PBS. Fixation was performed by adding 1 mL of cold 70% ethanol dropwise to the cell pellet while vortexing to ensure thorough fixation and minimize clumping. Cells were fixed for at least 30 mins on ice. Fixed cells were washed twice with PBS. To selectively stain DNA, cells were treated with 50 µL of RNase A, followed by the addition of 400 µL propidium iodide (PI) solution (50 µg/ml) per million cells directly to the RNase A-treated pellet. The samples were mixed thoroughly and incubated at room temperature for 10 mins. The stained samples were analysed by flow cytometry and data was collected from at least 20,000 single cells per condition. Data was plotted using floreada software (https://floreada.io).

### Immunoblotting

Cell pellets were resuspended in RIPA buffer, incubated for 20 mins on ice and then centrifuged at 13300 rpm for 20 mins at 4 °C. The supernatant was transferred to a fresh Eppendorf tube, and protein concentration was measured with a Bradford assay and adjusted to the same concentration. Samples were resuspended to 1x sample buffer (125 mM Tris–HCl pH 6.8, 10% Glycerol, 1% SDS, 0.2% (w/v) Orange G, 10% β-mercaptoethanol) and boiled at 95 °C for 5 mins. 10 µg of protein was loaded per lane in a 4-20% SDS-PAGE gel (BioRad), followed by transfer to a nitrocellulose membrane (BioRad Transblot Turbo). Membranes were blocked in 5% (w/v) Milk or BSA in TBS-T (20 mM Tris–HCl pH 7.5, 150 mM NaCl, 0.1% Tween-20) for 1 hr and incubated with primary antibodies at 4°C overnight with gentle rocking. Membranes were washed with TBS-T for 10 mins (x 3), followed by incubation with secondary antibody diluted in 5% (w/v) milk or BSA in TBS-T for 1 hr at room temperature with gentle rocking, followed by washing with TBS-T for 10 mins (x 3). For fluorescent antibodies, membranes were visualised with an Odyssey Fc Gel Imager (LI-COR). For HRP-conjugated secondaries (Fig S2B), ECL reagent (BioRad Clarity ECL Reagent 1705061) was used to detect, and membranes were visualised on X-ray films (Cytiva #GE28-9068-37), developed on an OPTIMAX 2010 X-ray Film Processor (PROTAC-Med).

### CUT&RUN Sequencing and Library Preparation

CENP-A, H3K9me3 and H3K27me3 CUT&RUN were performed natively with 5 × 10^5^ cells using CUTANA™ ChIC/CUT&RUN Kit (EpiCypher, SKU 14-1048), according to manufacturer’s protocol (version 2). The following antibodies and concentrations were used: CENP-A (Enzo ADI-KAM-CC006, 1:50), H3K9me3 (Abcam ab8898, 1:50) and H3K27me3 (C36B11, 1:50). DNA sequencing libraries were prepared using the NEBNext Ultra II DNA Library Prep Kit for Illumina (NEB) following published protocol. For multiplexing, we used NEBNext Multiplex Oligos for Illumina (Index Primers Set 1 and 2). Size selection was performed using Ampure XP beads (Beckman Coulter) to isolate mononucleosomal DNA fragments of approximately 150-180 bp (excluding adapters). Library yield and quality were evaluated using the Qubit HS dsDNA Quantification Assay Kit (Thermo Fisher Scientific) and the TapeStation 4150 System (Agilent). Multiplexed libraries were diluted to final concentrations of 1, 2, or 4 nM and sequenced at 1.8 pM on a NextSeq 500 system (Illumina) with the NextSeq 500/550 High Output v2.5 (75 cycles) kit (Illumina).

For the analysis of CUT&RUN data, raw FASTQ files were downloaded and concatenated per sample using basic Unix commands. Read quality was assessed using Basespace (Illumina) and FastQC software^71^. Reads were then aligned to the RPE1 genome using Bowtie2^72^, with trimming parameters adjusted based on read quality. SAM to BAM file conversion, BAM sorting, and indexing were carried out using SAMtools v1.16^73^. Duplicate reads were removed using the MarkDuplicates command in Picard^74^. The resulting sorted and duplicate-removed BAM files were used to calculate read counts, normalize data (counts per million, CPM), and convert to BigWig format using the bamCoverage command in Deeptools v2^75^. BigWig files were visualized using IGV and with pyGenomeTracks^76^. Read count matrices were generated using the multiBigWigSummary command in Deeptools v2 and output visualised using matplotlib.

### MBP-nanobody-Hia5-His6 Purification

Mouse and Rabbit MBP-nanobody-Hia5-6xHis constructs were a gift from Aaron Straight (Stanford University). MBP-nanobody-Hia5 proteins were overexpressed by rhamnose induction in BL21 DE3 cells for 16-20 hours at 18 degrees before harvesting by centrifugation. Cells were pelleted and resuspended in Lysis Buffer (50 mM HEPES, pH 7.5; 300 mM NaCl; 10% glycerol; 0.5% Triton X-100) with 1 ETDA-free protease inhibitor tablet per litre of culture. The cells were then lysed by probe sonication (Qsonica Q125) (6 pulses, 30s on, 1 min off at 200W). After a clearance centrifugation at 40,000 x g for 1 hour, the supernatant was added to washed Ni-NTA beads and mixed for 1 hour at 4 degrees. The slurry was added to a gravity flow column, washed with 50 mM HEPES, pH 7.5; 300 mM NaCl followed by 50 mM HEPES, pH 7.5; 300 mM NaCl; 20 mM imidazole. The protein was eluted by 50 mM HEPES, pH 7.5; 300 mM NaCl; 250 mM imidazole in 30ml, confirmed by SDS-PAGE. The elution was then dialysed to remove the imidazole at 4 degrees overnight in 50 mM Tris-HCl, pH 8.0; 100 mM NaCl; 1mM DTT. After dialysis, the protein was concentrated, filtered and applied to a Recourse Q anion exchange column. After confirmation of protein purity by SDS-PAGE, the protein was concentrated to 1mg/ml, snap frozen and stored at −80 degrees until use.

### CENP-A and H3K9me3 DiMelo-seq

DiMeLo-seq was performed according to the standard protocol in Maslan et al^77^ with some minor changes. All buffer compositions are as per published protocol^77^. Briefly, 6×10^6^ RPE cells were pelleted and resuspended in 1x PBS. For IAA experiments, cells were fixed with 0.1% paraformaldehyde (PFA) by adding 6.2 µl of 16% PFA to 1 ml of cell suspension with gentle vortexing for 2 mins. Fixation was quenched with 1.25 M glycine (60 µl per 1 ml of suspension), followed by centrifugation at 500 × g for 3 mins at 4°C. The supernatant was removed, and the fixed cells were resuspended in Dig-Wash buffer (0.02% Digitonin).

For all other experiments (native DiMelo-seq), 6×10^6^ RPE cells were washed in 1x PBS and pelleted at 600 x g. Supernatant was removed, and the cell pellet was resuspended in 1 mL Dig-Wash buffer and incubated on ice for 5 mins. Nuclei were centrifuged at 600 x g and the supernatant was removed. Permeabilization quality was assessed using Trypan Blue staining. Nuclei pellets were gently resuspended using wide-bore pipette tips in 400 µl Tween-Wash buffer containing the primary antibody (CENP-A, 1:50 or H3K9me3, 1:50). Samples were incubated on a hula mixer at 4°C overnight. After incubation, samples were centrifuged, supernatant removed and washed twice with 0.95 mL Tween-Wash buffer. Each wash involved thorough resuspension of the pellet with pipetting, followed by rotation at 4°C for 5 minutes before centrifugation.

To quantify the MBP-Nb-Hia5 protein, frozen protein aliquots were thawed at room temperature and immediately placed on ice. Concentration was measured using a Qubit fluorometer with a 2 µl sample volume. Nuclei pellets were resuspended in 400 µl Tween-Wash buffer containing exactly 200 nM pA-Hia5. Binding was performed on a hula mixer at 4°C for 2 hrs. After incubation, samples were centrifuged, supernatant removed and washed twice with 0.95 ml Tween-Wash buffer as described above. Nuclei pellets were then resuspended in 200 µl of Activation Buffer per sample, supplemented with 800 µM S-adenosylmethionine (SAM) as a substrate for the Hia5 methyltransferase. Samples were mixed every 30 mins and SAM was replenished after 1 hr. After 2 hr SAM incubation, nuclei were centrifuged, supernatant removed and resuspended in 40 µl of 1x PBS.

### UHMW DNA Extraction and Library Preparation

High-molecular-weight (HMW) genomic DNA was extracted using the NEB Monarch® HMW DNA Extraction Kit for Tissue. The 40 µl of resuspended nuclei were moved to a 2 ml Eppendorf DNA LoBind tube. Nuclei were lysed by adding 1.8 ml of NEB Monarch® HMW gDNA Tissue Lysis Buffer premixed with 60 µl of proteinase K. The mixture was gently pipetted 10 times with a wide-bore pipette and incubated at 56°C for 10 minutes. RNase A (15 µl) was then added, followed by gentle mixing and incubation at 56°C for an additional 10 minutes with agitation at 650 rpm on a heat block. DNA was precipitated in a 5 mL Eppendorf DNA LoBind tube by adding 2.5 ml of isopropanol with three glass beads to the lysed nuclei and rotating at 10 rpm for 20 minutes. After resting at room temperature, isopropanol was aspirated, and the DNA adsorbed to beads was washed twice with 2 ml of Wash Buffer. The DNA-bead complex was transferred to a bead retainer, and residual Wash Buffer was removed by a soft spin. Beads were then immediately transferred to 2 ml Eppendorf tubes containing 300 µl of Extraction EB (EEB, ONT SQK-ULK114 kit) to elute the DNA. Beads were incubated in EEB overnight at room temperature. The eluate was separated from the beads by centrifugation and adjusted to a final volume of 750 µl with additional EEB. This 750 µl volume was then taken forward to the tagmentation step as per the standard SQK-ULK114 protocol. CENP-A and H3K9me3 DiMeLo-seq DNA was sequenced on a PromethION 2 Solo device (ONT) with R10.4.1 flow cells with adaptive sampling for all centromeres and all of chromosome 4, to a depth of 25 Gb (CENP-A) and 20 Gb (H3K9me3).

### Data Processing of DiMelo-seq

Pod5 files were basecalled using Dorado v0.7.3 using dorado basecaller <u>sup@v4.3.0</u> for 6mA and 5mCG/5hmCG (https://github.com/nanoporetech/dorado). Basecalled bam files were then aligned to the RPE1 genome^39^ using dorado. Aligned bam files were then sorted using samtools (v1.16.1), filtered with samtools view -F 2308 to remove unmapped (0×4) non-primary alignments (0×100) and supplementary alignment (0×800), following by indexing. Indexed bam files were then processed through a refModMatch script (https://github.com/altemoselab/miscTools/blob/main/refModMatch.py) to ensure we only keep bases in the MM tag if they are a match to the reference genome.

6mA and 5mCG modification filtering was performed using modkit (https://github.com/nanoporetech/modkit) using modkit pileup with the following thresholds: -- filter-threshold A:0.8 --mod-threshold a:0.98 --filter-threshold C:0.8 --mod-threshold m:0.8. 6mA thresholds were tuned for specificity and signal-to-noise by using endogenous CENP-A auxin depletion (Fig S5) where endogenous CENP-A is fully depleted. Bigwigs for visualisation were generated from columns 1, 2, 3 and 11 (fraction modified) of the output bedmethyl file from Modkit. To visualise single molecules, we used fibertools-rs (v0.5.4; https://github.com/fiberseq/fibertools-rs)^78^, and used the ft extract tool to extract --m6A and -- cpg motifs into a bed file with --ml 250 and --ml 204 thresholds respectively for each modification (in line with modkit filtering above). Bigwig files and single molecule tracks were visualised with pyGenomeTracks^76^.

### Centromere Annotation, Identification of HORs and CDRs

Alpha satellite annotations were previously identified using HumAS-HMMER For AnVIL (https://github.com/fedorrik/HumAS-HMMER_for_AnVIL)^39^. The resulting annotations were compared with CHM13 CenSat annotations^3^ to identify active, inactive and divergent HORs, as well as monomeric satellites in the RPE1 genome. For analysis of active HORs in Fig 4 and Fig 5, active HOR annotations were lifted to include only the CDR/CENP-A-containing active HOR. Cases such as Cen3 (Hap1 and Hap2), Cen4 (Hap1), and Cen13 (Hap2) which contain multiple split active HORs were contained to just the CENP-A-containing HOR. These CENP-A-containing active HORs were normalised to the same length (see below) and plotted as active HOR +/− 1 Mb, with flanking regions broadly defined as “pericentromere”.

### Normalisation of Active HOR and CDR length

To standardize regions of varying lengths for heatmap analysis, each region was normalized to a fixed number of bins. Regions were extended symmetrically with upstream and downstream flanking areas. For Active HORs, regions were normalized to 1000 bins, with flanking pericentromeric regions (±1 Mb) divided into 500 bins each. CDRs were scaled to 200 bins, while each flank (±125 kb) was divided into 125 bins. Mean signal intensities for each bin were extracted from BigWig files using the pyBigWig.stats() function, with missing values replaced by NaN. The normalized signal profiles for all regions were combined into matrices for visualization, where each row represents a single Active HOR or CDR, sorted numerically (e.g., chr1_hap1, chr1_hap2, chr2_hap1, etc.).

### CDR Length Analysis and New CDR Detection

CDRs were initially identified with CDR-Finder (https://github.com/EichlerLab/CDR-Finder)^58^ using the following parameters: height_perc_valley_threshold: 0.39; prom_perc_valley_threshold: 0.45. All other parameters were as per default test parameters. After initial CDR detection, each CDR bed file was manually inspected for accuracy and compared to CENP-A DiMelo-seq data. Any non-CENP-A containing CDRs were manually removed. CDRs for Chr7 Hap1 and Chr10 Hap2 were manually annotated for accuracy. To ensure comparison of only primary CDRs, all non-primary/newly-formed CDRs were removed. The total length of each CDR was then summed and directly compared on a per centromere basis to the corresponding parent centromere. To detect newly-formed CDRs, CDR-Finder^58^ output was manually inspected as above for accuracy, without removing non-primary CDRs. All CDRs within 500kb were merged and considered as one single centromere locus. Any CENP-A-containing CDRs not present in the parent CDR database were considered newly formed i.e. any new centromere peak more than 500 kb from the primary CDR site. The following chromosomes were not considered in all CDR analyses: Cen4 Hap2 (neocentromere-containing haplotype); CenX Hap2 (not present in genome), Cen18 Hap1 and Cen18 Hap2 (poorly-defined CDRs).

### Mitotic Spreads

Cells were treated with KaryoMAX™ Colcemid™ (100 ng/ml, Gibco) for 3 hrs and mitotic cells were collected by mitotic shake-off. To prepare cells for spreading, a 75 mM KCl solution was prewarmed in a 37 °C water bath, and a fresh Carnoy fixative solution (methanol/acetic acid, 3:1) was prepared and placed on ice. The cell suspension was centrifuged at 500 × g for 5 minutes, and the supernatant was discarded, leaving a small residual volume in which the pellet was gently resuspended. For hypotonic treatment, 1 mL of 75 mM KCl (prewarmed to 37 °C) was slowly added to the cell suspension, followed by an additional 4 mL of the same solution. The tubes were incubated in a 37 °C water bath for 15 mins, with gentle shaking once or twice during incubation to homogenize the solution. After incubation, the cells were centrifuged at 500 × g for 5 minutes at 4 °C. The supernatant was removed, leaving approximately 500 μL with the cell pellet, and the tubes were immediately placed on ice.

To fix the cells, the pellet was gently resuspended in the residual supernatant. While the tube was vortexed at the lowest speed, 1000 μL of ice-cold fixative was added dropwise, followed by an additional 10 mL of fixative. The suspension was centrifuged at 230 × g for 10 minutes at 4 °C, and the cell pellet was resuspended in 10 mL of fresh fixative. The prepared cells were stored at −20 °C until ready for chromosomal spreading.

For dropping onto slides, Epredia Superfrost Plus slides were cleaned and kept in a coplin jar with methanol and kept at 4 °C until required. When ready for dropping, a cleaned slide is removed and placed tilted on an Eppendorf tube rack and 100 μL of fixed cells were dropped from a ∼30 cm height onto the slide and allowed to flow down the glass. Excess fixative was removed on a paper towel and the slide was allowed to dry fully for ∼5 mins.

### Immunofluorescence

Immunofluorescence on methanol-acetic acid fixed cells was performed as described in Beh et al. (2016)^79^ with some modifications. The primary antibody used to identify the dicentric chromosome was a guinea pig anti-CENP-C (1:1000, MBL, PD030) and the secondary antibody was used at 1:500 (anti-guinea pig Alexa Fluor® 488, Jackson ImmunoResearch Laboratories). Imaging was performed on a Thunder Live Cell (Leica) equipped with a Leica K8 CMOS Back-thinned camera (95% quantum efficiency). Images ≤4 μm thick were captured in 0.2 μm z-sections at room temperature using a 100x Leica HCX PL APO oil-immersion objective (numerical aperture 1.4), and then projected and post-processed (Instant Computational Clearing) using <u>Leica LAS-X</u> software.

### Multicolor-FISH Karyotyping (M-FISH)

For M-FISH following immunocytochemistry, slides were washed in PBS for 3 min as described in Beh et al. (2016)^79^ and Multicolor FISH was performed with 24 XCyte probe (MetaSystems) following manufacturer’s instructions. The Metafer imaging platform (MetaSystems, Metafer4 v3.13.5) and the Isis software (v5.8.14) were used for automated acquisition of the chromosome spread and M-FISH images analysis.

## Acknowledgements

We thank members of the Jansen lab, past and present for discussion and critical analysis of this manuscript. We thank Luke Fulcher for technical advice regarding lentiviral targeting, Aaron Straight for providing early access to the nanobody-Hia5 constructs, Cy Chittenden from the Altemose lab for advice on Hia5 purification, Karen Miga and Brandy McNulty for advice on UHMW DNA extraction. We further thank Robert Hedley and Vasiliki Tsioligka for providing technical assistance in cell cycle analysis and single-cell sorting at The Don Mason Facility of Flow Cytometry, Sir William Dunn School of Pathology, University of Oxford. We also acknowledge the Cell and Tissue Imaging Platform (PICT-IBiSA, a member of the French national research infrastructure France-BioImaging; ANR10-INBS-04). This work was funded by a UKRI Marie Sklodowska**-**Curie Actions European Postdoctoral Fellowship (101066059), a Wellcome Early Career Award (303812/Z/23/Z) and John Fell Award (0013250) (Oxford University Press) to BLC. DF receives salary support from the CNRS, MD is supported by Institut Curie. Experimental work in the Fachinetti lab is supported by ANR-21-CE13-0030. SG receives salary support from the Rita Levi Montalcini Award, EV is supported by a PNRR Ph.D. Fellowship from the Italian Ministry of University and Research (MUR), and work in the Giunta lab is supported by AIRC Start-Up grant 202 ID. # 25189, Sapienza Excellence grant and ERC CENTROFUN grant # 101078838 to SG. Work in the Altemose Lab was supported by a Howard Hughes Medical Institute Hanna H. Gray Faculty Fellowship and by the Chan Zuckerberg Biohub. LETJ and the Jansen lab are generously funded by a Wellcome Senior Research Fellowship 210645/Z/18/Z and a Wellcome Discovery Award UNS53865.

## Supplemental Figures

**Figure S1:**
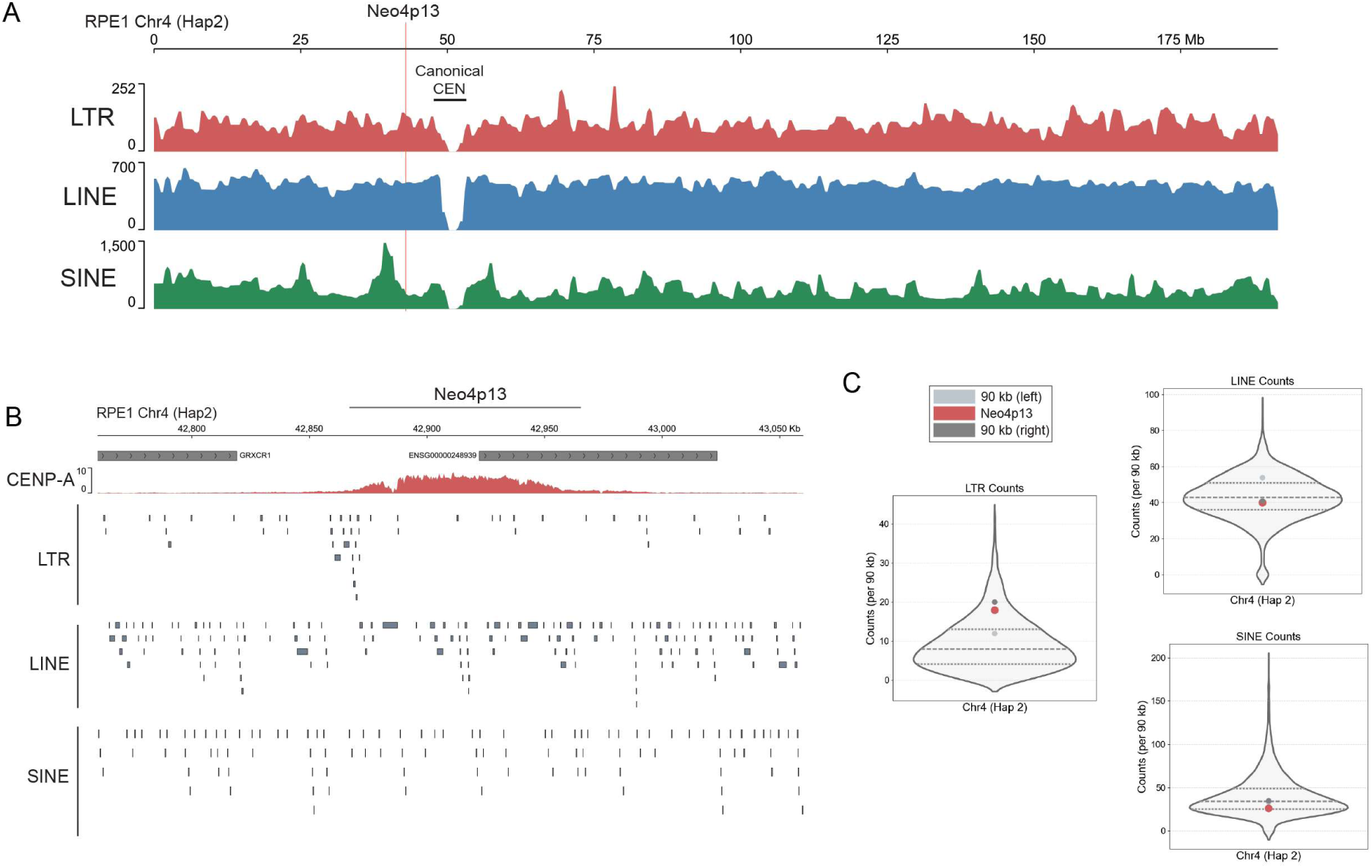
Transposable Element Density across Chromosome 4. **(A)** Density plot showing the distribution of the three main classes of retrotransposable elements (TEs) along chromosome 4 (LTRs, LINES and SINES) analyzed in 1 Mb intervals. A distinct dip in TE density is observed within the centromeric regions, reflecting their characteristic satellite composition. **(B)** Retrotransposon density across the Neo4p13 domain in 90 kb bins on chr4 (hap2). **(C)** Violin plot of total LTR, LINE and SINE density across entire chr4 (hap2), highlighting bins directly overlapping with or immediately adjacent of Neo4p13.

**Figure S2:**
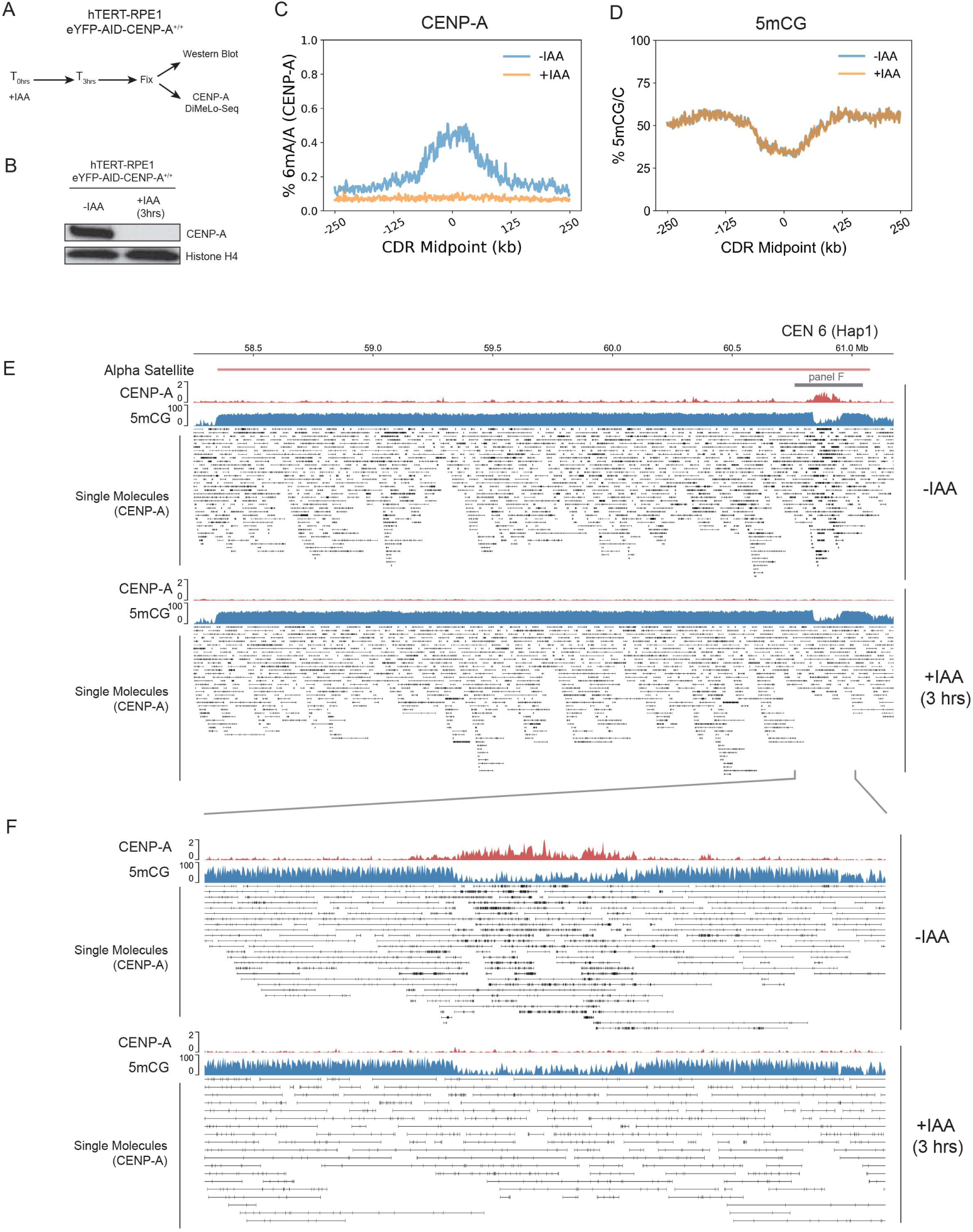
On-target specificity of CENP-A-DiMelo-seq. **(A)** Schematic of 3 hr auxin depletion using hTERT-RPE1 eYFP-AID-CENP-A^+/+^ cells^51^. **(B)** Immunoblot for CENP-A and Histone H4 on nuclear extracts +/− 3hr IAA treatment. **(C)** CENP-A-directed 6mA/A plot with the mean 6mA trace for +/− IAA conditions for all centromeres, overlayed for comparison. **(D)** Combined 5mCG/C plot with the mean 6mCG trace for +/− IAA conditions for all centromeres, overlayed for comparison. **(E)** View of the entire HOR of CEN6 (Hap1) +/− IAA conditions with CENP-A 6mA/A and 5mCG/C averaged genomic tracks, and single molecule alignments with CENP-A-directed 6mA modifications. **(F)** Zoom into the CDR of CEN6 (Hap1) +/− IAA conditions with CENP-A 6mA/A and 5mCG/C averaged genomic tracks, and single molecule alignments with CENP-A-directed 6mA modifications.

**Figure S3:**
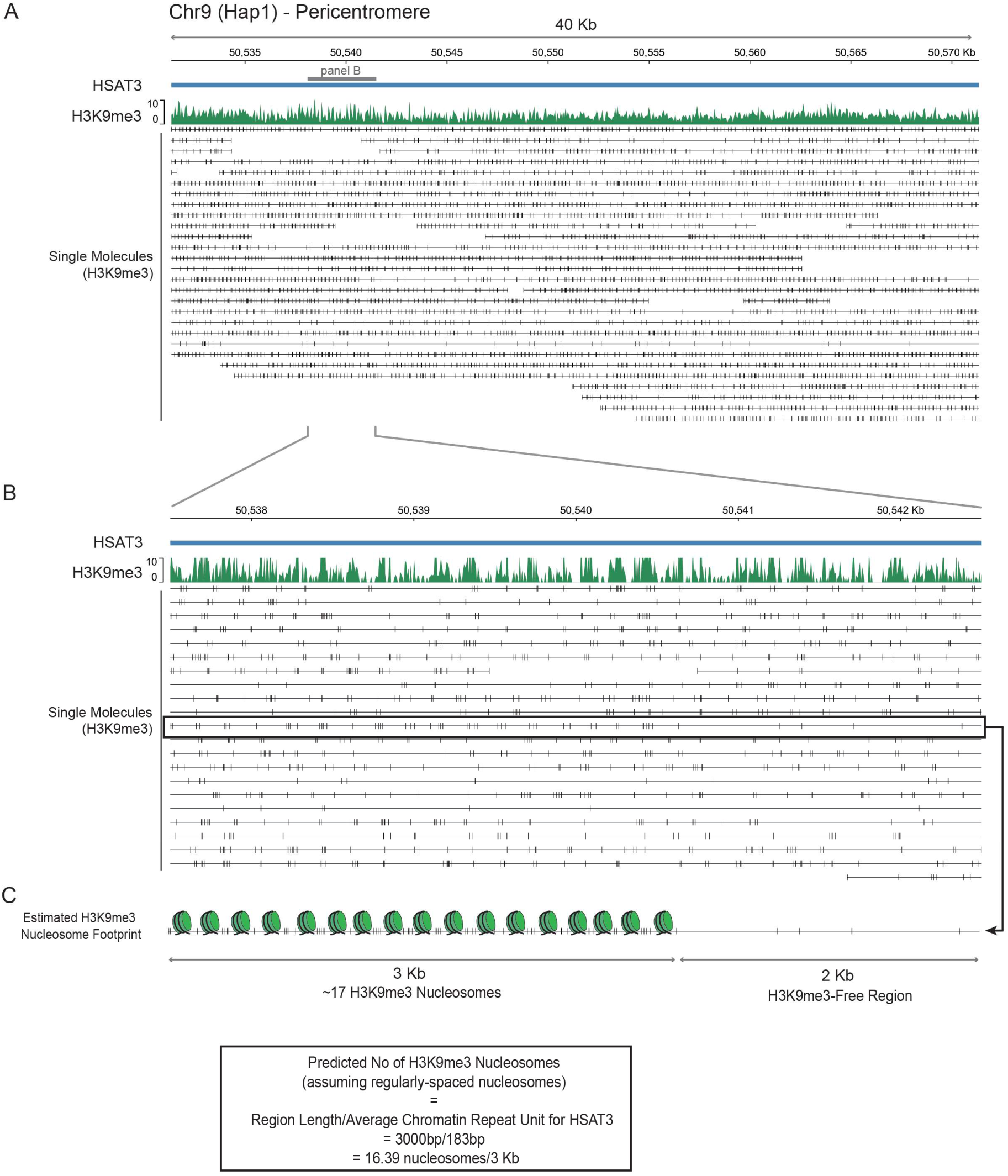
Nucleosome-level sensitivity of H3K9me3 DiMelo-seq at constitutive pericentromeric heterochromatin. **(A)** Visualisation of a 40 kb region on human satellite 3 (HSat3) of chromosome 9, showing H3K9me3 averaged m6A/A track (green) and single molecule alignments (black, bottom). Black vertical lines represent H3K9me3-directed 6mA modification. **(B)** Zoom in to a 5 kb region of HSat3 showing H3K9me3 averaged m6A/A track (green) and single molecule alignments (black, bottom). 6mA modifications appear as regularly spaced nucleosome-like intervals. **(C)** Estimation of H3K9me3 nucleosome footprinting on a 5kb region of HSat3, showing a 3 kb H3K9me3-dense and a 2 kb H3K9me3-free region. On this single molecule, we can estimate nucleosome positioning of H3K9me3-containing nucleosomes to be approximately 17 nucleosomes within this 3kb H3K9me3-dense region (176 bp/nucleosome, which is consistent with previous estimates of 183 bp/nucleosome for HSat3 heterochromatic regions based on Fiber-Seq measurements^55^.

**Figure S4:**
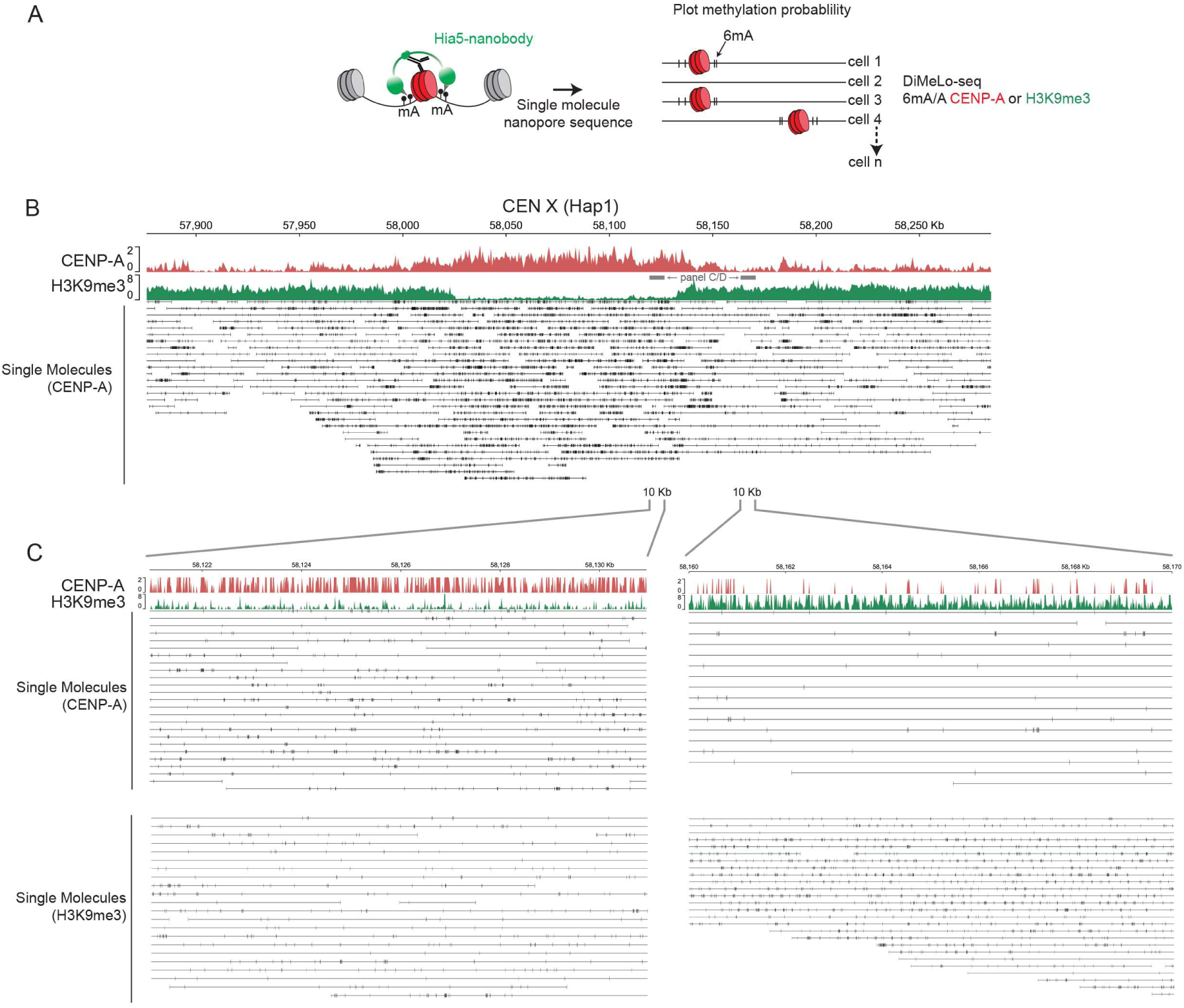
Nucleosome-level view of CENP-A and H3K9me3 DiMelo-seq at centromere boundary. **(A)** Schematic of DiMelo-Seq workflow and visualisation of single molecule alignment. Black vertical lines represent 6mA modification. **(B)** Centromere X (Haplotype 1) visualised with CENP-A (red, 6mA/A) and H3K9me3 (green, 6mA/A). Single-molecule alignments with 6mA modifications for CENP-A (bottom), show the specificity of the CENP-A signal to the CDR. **(C)** 10kb zoom in to regions inside (left) versus outside (right) of the CENX CDR for single DNA molecules, showing the specificity for CENP-A and H3K9me3 detection at our chosen m6A probability filtering.

**Figure S5:**
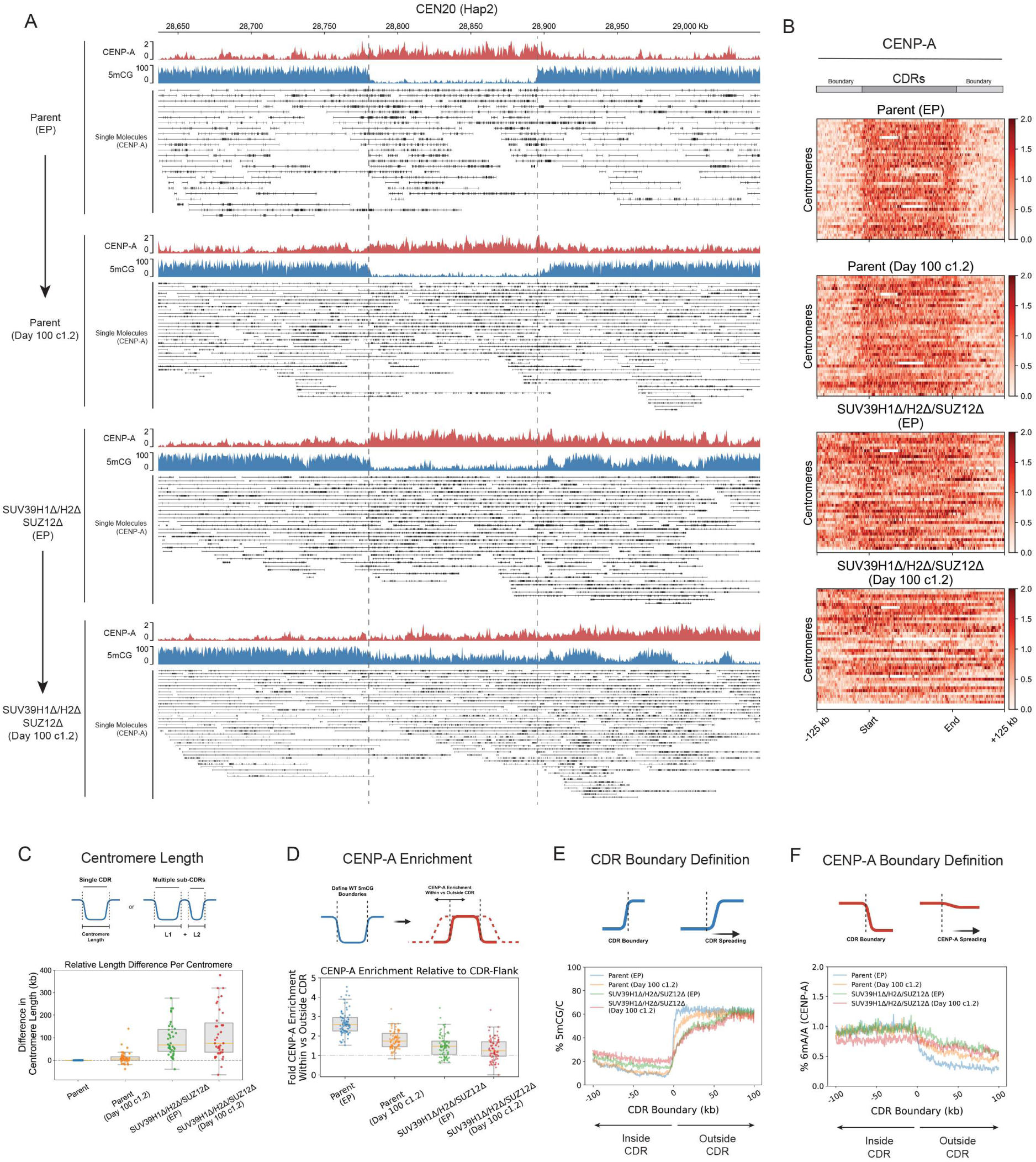
Canonical centromere drift over time, restricted by heterochromatin. **(A)** View of the entire HOR of CEN 20 (Hap 2) for Parental (RPE-Neo4p13), Parental (Day 100 Cone 1.2), SUV39H1Δ/H2Δ/SUZ12Δ (early Passage, EP) and SUV39H1Δ/H2Δ/SUZ12Δ (Day 100 Clone 1.2) with aggregate DiMelo-Seq enrichments for CENP-A (red) and 5mCG enrichment (blue). Single molecules alignments for CENP-A-directed 6mA (black). **(B)** Heatmaps for each mutant illustrating the distribution of CENP-A (red) across Centromere Dip Regions (CDRs), as defined by the boundary of 5mCG dips on either side of each CENP-A-containing region in WT RPE-Neo4p13. Each line on the heatmap represents a single CDR/Centromere. All CDRs have been sorted numerically and scaled to the same length for comparison. **(C) Centromere length** as defined by the difference in sum total length of CDR/sub-CDRs relative to the corresponding parent centromere. Box plots summarize the distribution for parent and mutant lines, while individual data points are shown as coloured dots. A dashed reference line at 1 indicates early passage parental centromere length. The orange line represents the median centromere length. **(D) CENP-A Enrichment:** Box plot depicting the ratio of CENP-A enrichment within CDRs relative to their flanking regions (±100 Kb) for parent and mutant lines. Individual data points are shown as coloured dots, while box plots represent the distribution of ratios. A dashed reference line at 1 indicates equal enrichment within and outside the HORs. The orange line represents the median CENP-A ratio. **(E) CDR boundary definition:** plot of all 5mCG CDR boundary profiles centred on CDR boundaries (±100 Kb). Mean of combined aggregated traces is shown for all samples. **(F) CENP-A boundary definition:** Plot of CENP-A enrichment inside versus outside the CDR boundary (defined by Parental 5mCG) for each cell line (±100 Kb). Mean of combined CENP-A (6mA/A) aggregated trace is shown for all samples.

**Figure S6:**
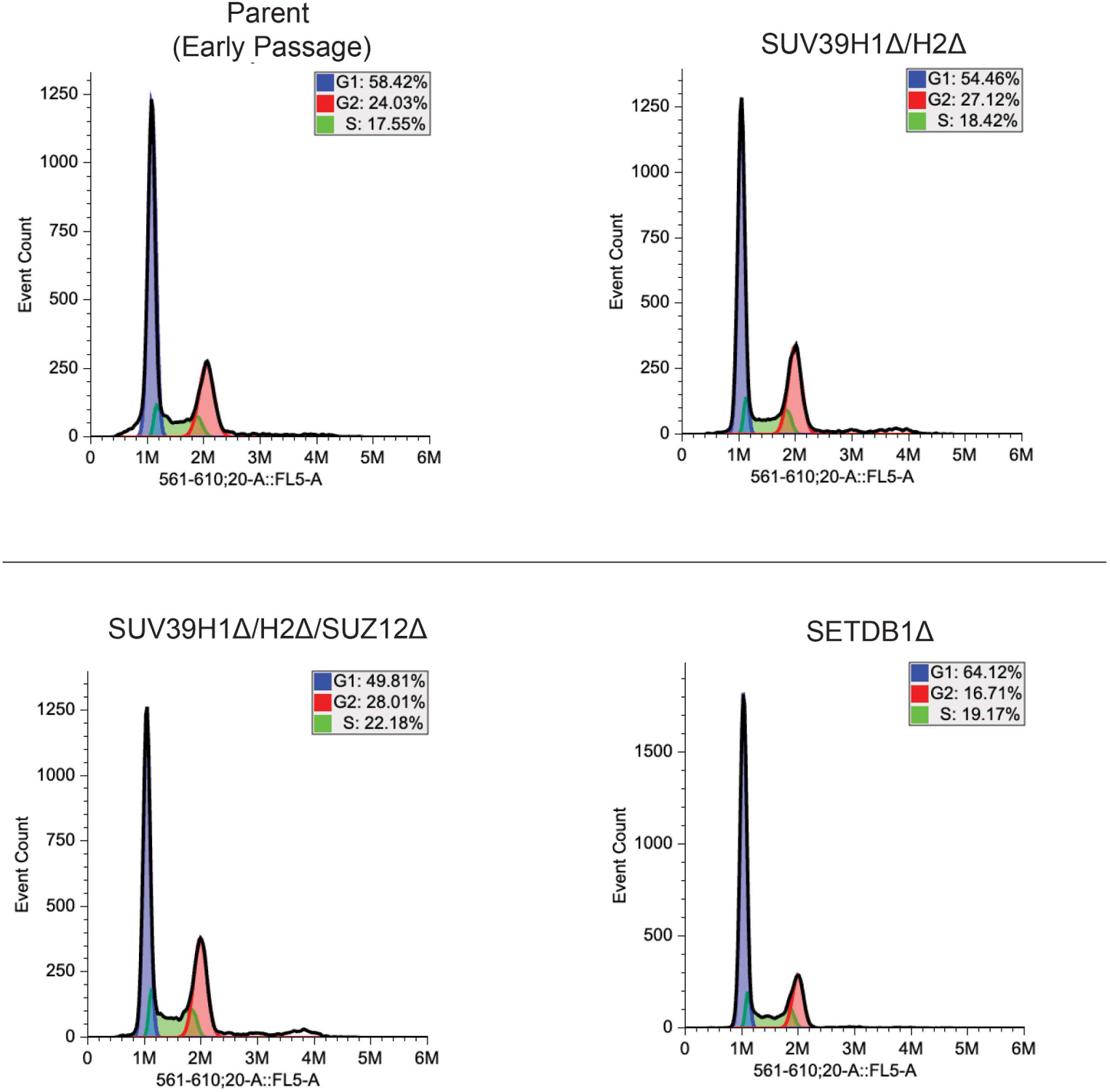
Cell cycle profiles, ploidy and karyotypes of cell lines. Measure of cell cycle distribution and ploidy by Propidium Iodide DNA-staining and flow cytometry for each cell line used in this study.

**Figure S7:**
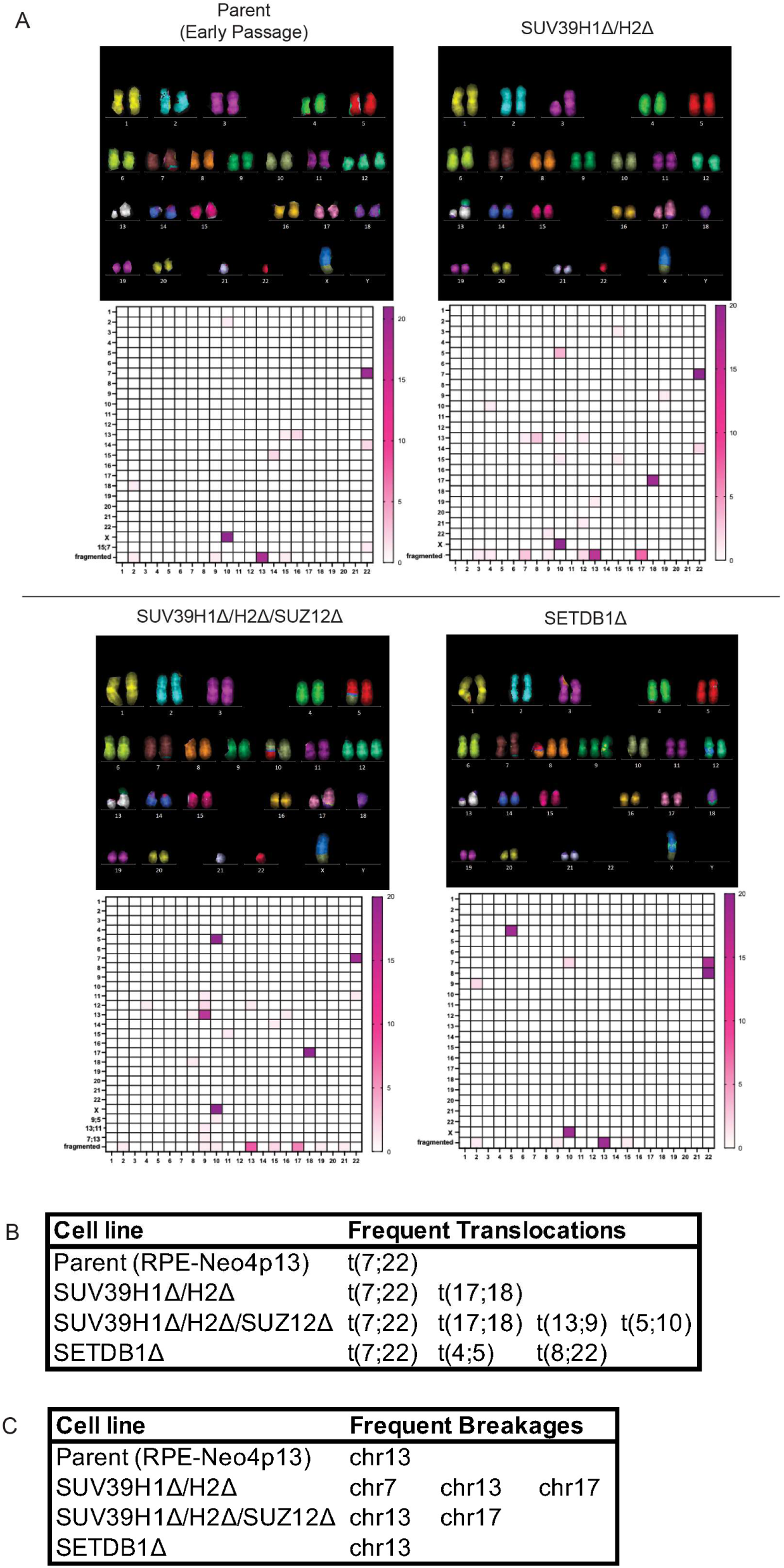
Ploidy and karyotypes of cell lines. **(A)** Representative mFISH karyotypes for each cell line used in this study, alongside heatmap (below) indicating the frequency of translocations and chromosome breakages for each cell line based on 20 mFISH images per cell line. **(B)** Table listing most frequent translocations identified in each cell line. **(C)** Table listing most frequent chromosome breakages identified in each cell line.

## Supplemental Tables

**Table 1:** Summary of the coordinates of all new CENP-A containing CDRs.

| Cell Line | Chromosome | Start | End |
| --- | --- | --- | --- |
| Parent 100d Clone 1.2 | Chr21 (Hap1) | 4830300 | 4895300 |
| SUV39H1Δ/H2Δ/SUZ12Δ | Chr1 (Hap1) | 125004534 | 125149534 |
| SUV39H1Δ/H2Δ/SUZ12Δ | Chr1 (Hap1) | 126269534 | 126354534 |
| SUV39H1Δ/H2Δ/SUZ12Δ | Chr9 (Hap2) | 46215681 | 46365681 |
| SUV39H1Δ/H2Δ/SUZ12Δ | Chr20 (Hap1) | 27068226 | 27113226 |
| SUV39H1Δ/H2Δ/SUZ12Δ | Chr21 (Hap1) | 4650300 | 4860300 |
| SUV39H1Δ/H2Δ/SUZ12Δ 100d Clone 1.2 | Chr1 (Hap1) | 125109534 | 125149534 |
| SUV39H1Δ/H2Δ/SUZ12Δ 100d Clone 1.2 | Chr1 (Hap1) | 126269534 | 126419534 |
| SUV39H1Δ/H2Δ/SUZ12Δ 100d Clone 1.2 | Chr3 (Hap2) | 96531917 | 96561917 |
| SUV39H1Δ/H2Δ/SUZ12Δ 100d Clone 1.2 | Chr9 (Hap2) | 46215681 | 46360681 |
| SUV39H1Δ/H2Δ/SUZ12Δ 100d Clone 1.2 | Chr20 (Hap1) | 27063226 | 27128226 |
| SUV39H1Δ/H2Δ/SUZ12Δ 100d Clone 1.2 | Chr21 (Hap1) | 4645300 | 4865300 |
| SETDB1Δ | Chr7 (Hap1) | 65533216 | 65578216 |
| SETDB1Δ | Chr9 (Hap1) | 45447340 | 45487340 |
| SETDB1Δ | Chr19 (Hap2) | 29111633 | 29296633 |
| SETDB1Δ | Chr20 (Hap2) | 27113480 | 27193480 |
| SETDB1Δ | Chr21 (Hap1) | 4825300 | 4895300 |

**Table 2:**
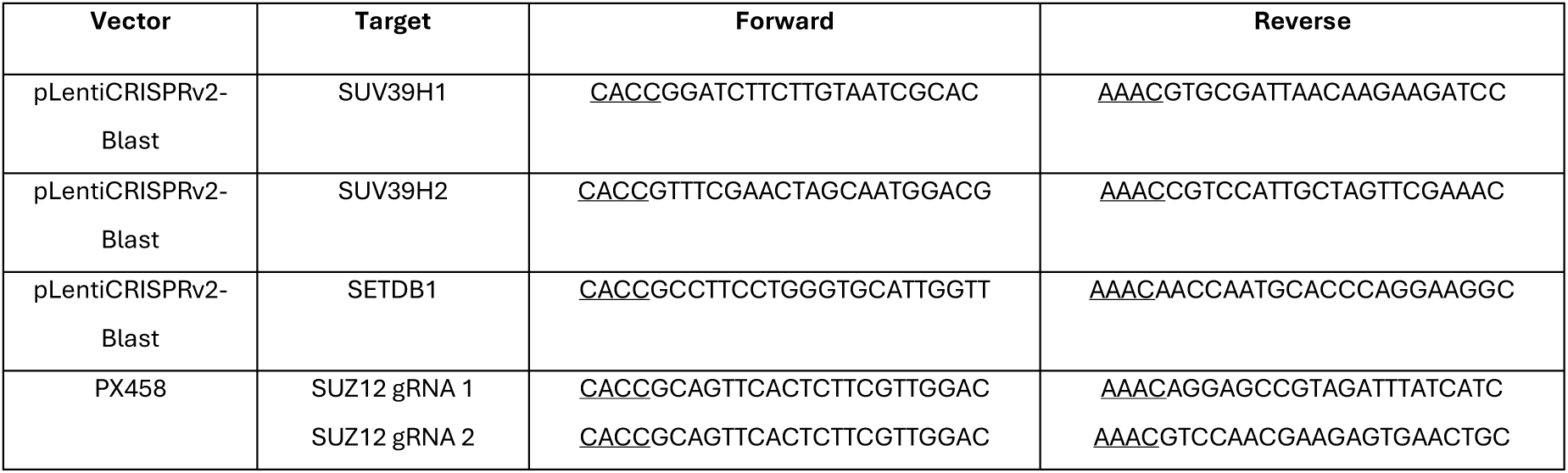
Table of gRNA Primers used for CRISPR knockouts.

**Table 3:** Primary and Secondary Antibodies used for Immunoblotting.

| Target | Antibody | Working dilution |
| --- | --- | --- |
| SUV39H1 | Cell Signalling (D11B6) | 1:1000 |
| SUV39H2 | Abcam (ab190870) | 1:1000 |
| SUZ12 | Cell Signalling (D39F6) | 1:1000 |
| SETDB1 | Cell Signalling (C1C12) | 1:1000 |
| CENP-A | Cell Signalling (2186S) | 1:1000 |
| Histone H4 | Merck Millipore (05-858) | 1:1000 |
| α-tubulin | Sigma (T9026) | 1:5000 |
| H3K9me3 | Abcam (ab8898) | 1:1000 |
| H3K27me3 | Cell Signalling (C36B11) | 1:1000 |
| Rabbit IgG | IRDye® 800CW (926-32211) | 1:5000 |
| Mouse IgG | Rockland DyLight™ 680 (610-744-124) | 1:5000 |
| Rabbit IgG | Jackson ImmunoResearch HRP (111-035-008) | 1:5000 |

## Notes

### Competing Interest Statement

The authors have declared no competing interest.

